# Rapid disassembly and Piwi-independent reassembly of *Drosophila* piRNA cluster heterochromatin following acute heat shock

**DOI:** 10.1101/2023.07.26.549158

**Authors:** Nicholas Rice, Samantha Ho, Zhiping Weng, William E. Theurkauf

## Abstract

Environmental stress activates transposons and is proposed to generate genetic diversity that facilitates adaptive evolution. piRNAs guide germline transposon silencing, but the impact of stress on the piRNA pathway is not well understood. In *Drosophila,* the Rhino-Deadlock-Cuff complex (RDC) drives transcription of clusters composed of nested transposon fragments, generating precursors that are processed into mature piRNAs in the cytoplasm. We show that acute heat shock triggers rapid, reversable, loss of RDC localization and cluster transcript expression with coordinate changes in the cytoplasmic processing machinery. Maternal piRNAs bound to Piwi are proposed to guide Rhino localization to clusters during early embryogenesis. However, RDC re-localization after heat shock is accelerated in *piwi* mutants and delayed in *thoc7* mutants, which disrupt piRNA precursor binding to THO complex, and we show that maternally deposited piRNAs are dispensable for RDC localization to the major 42AB cluster. Cluster specification is reconsidered in light of these findings.

## Introduction

### Environmental stress activates transposons in evolutionarily diverse systems

Although transposable element mobilization can induce deleterious mutations linked to disease (Payer and Burns 2019; Belancio, et al. 2009), McClintock proposed that stress induced transposon mobilization could generate beneficial genetic diversity, facilitating adaptive evolution (McClintock 1984). It is now clear that transposons have been co-opted for many essential host functions (Feschotte and Pritham 2007). However, transposition must occur in the germline to alter the inherited genome and drive evolution, and the germline response to stress is poorly understood (Mombach, et al. 2022; Kelleher, et al. 2020; Cappucci et al. 2019).

piRNAs guide transposon silencing in the germline, and this pathway is best understood in the *Drosophila* female germline. In this system, the 23 to 30 nucleotide small silencing piRNAs are derived from genomic loci composed of nested transposon fragments called clusters and from a subset of isolated transposon insertions bound by the HP1 homolog Rhino (Rhi) (Brennecke et al. 2007; Klattenhoff et al. 2009; Mohn et al. 2014; Shpiz et al. 2014)(Add Bergman Genome Biology, PMID: 17134480). The RDC complex, consisting of Rhino, Deadlock, and Cuff, drives transcription from both genomic strands, producing long un-spliced transcripts that are bound by the conserved THO complex and UAP56, which are components of the <u>Tr</u>anscription and <u>Ex</u>port (TREX) complex (Chen et al. 2016; Hur et al. 2016; Hur and Chung 2016; Mohn et al. 2014; Zhang et al. 2018; Zhang, Wang, et al. 2014). piRNA cluster transcripts are transferred from TREX to a specialized noncanonical export system that includes the *Drosophila* specific Nxf3 protein and CRM1 (*D. melanogaster* gene *embargoed*) (ElMaghraby et al. 2019; Kneuss et al. 2019; Mendel and Pillai 2019). Following export, these transcripts undergo ping-pong and phased processing. The ping-pong machinery localizes to perinuclear nuage granules and includes multiple Tudor domain proteins, the DEAD box RNA-binding protein Vasa, and the PIWI clade Argonaute proteins Aubergine (Aub) and Ago3 (Brennecke et al. 2007; Lim and Kai 2007; Webster et al. 2015; Xiol et al. 2014). Within the ping pong cycle, piRNAs bound to Aub guide cleavage of transposon transcripts, leading to post-transcriptional silencing and production of precursors for piRNA that bind Ago3 and guide cluster transcript cleavage, generating the precursors for Aub bound piRNAs (Brennecke et al. 2007). The ping-pong cycle also generates the precursors for phased processing, which generates head-to-tail arrays of piRNAs bound to Piwi, the founding member of the PIWI clade (Han et al. 2015; Mohn, Handler, and Brennecke 2015; Wang et al. 2015). The putative helicase Armi localizes to nuage and mitochondria and is proposed to shuttle precursors between these two compartments and facilitate phased processing with the mitochondrial nuclease Zuc (Ge et al. 2019). piRNA binding by Piwi triggers localization to the nucleus, where the complexes mediate transcriptional silencing with the interaction partner Panoramix /Silencio (Y. Yu et al. 2015; Sienski et al. 2015).

The piRNA response to heat stress has not been extensively characterized, but “heavy” heat shock, triggered by cycling flies between 4°C and 37°C, is reported to upregulate some transposable element families and lead to Ago3 localization to lysosomes (Cappucci et al. 2019). By contrast, a short 30 min heat shock at 38°C has been reported to trigger a modest change in piRNA levels mapping to transposons, but the target transposons are not upregulated, and piRNA levels were not reduced for elements that were upregulated (Funikov et al. 2015). In addition, long term maintenance of flies at 30°C is reported to lower piRNA expression while increasing ping-pong processing of some piRNAs (Fast and Rosenkranz 2018). These studies suggest that heat stress alters germline transposon expression, but the impact of heat shock on organization and function of the piRNA biogenesis machinery has not been systematically analyzed.

Here we show that acute heat shock leads to rapid reversible disassembly of piRNA cluster chromatin, including loss of the RDC and associated reduction in cluster transcript expression. Time course studies indicate that loss of TREX localization follows RDC dissociation, and that TREX localization recovers after the RDC. We also show that heat stress leads to reversible localization of Vasa to the nucleus and Aub to mitochondria, which may reflect trapping of these factors at normally transient compartments in the biogenesis cycle.

Maternally deposited piRNAs bound to Piwi have been proposed to guide RDC localization to clusters during oogenesis, while the propagation of Rhino on clusters is proposed to be Piwi and piRNA independent (Akkouche et al. 2017; Brennecke et al. 2008). However, here we show that *piwi* mutations actually accelerate recovery of Rhino localization following heat shock. Significantly, we also show that genetically eliminating maternal piRNAs mapping the major piRNA cluster at 42AB has no significant impact on Rhino binding or piRNA production from a paternally-supplied copy of this cluster. Heat shock thus provides novel insights into piRNA cluster chromatin assembly and calls into question the function of piRNAs and Piwi in this process.

## Results

### Heat Shock disrupts piRNA cluster heterochromatin

Heat shock increases transposon expression in plant and animal somatic cells (Belgnaoui et al. 2006; Raje, et al. 2018; Grandbastien et al. 1997; Roquis et al. 2021; Takehira et al. 2021; Sun et al. 2020; Garrigues et al. 2019), and stress induced transposition could facilitate adaptive evolution by generating beneficial genetic diversity. However, adaptive mutations must be transmitted through the germline to impact evolutionary fitness, and in *Drosophila* the germline response to heat shock has not been extensively characterized (Cappucci et al. 2019; Fanti et al. 2017; Jardim et al. 2015). To define the effects of heat shock on transposon expression in the *Drosophila melanogaster* germline, we incubated *w^1^* flies at 37°C for 1 hour and sequenced ovarian long RNAs 1 hour, 12hrs, and 24hrs after shifting back to room temperature. At 1hr post-heat shock, R1_DM, Copia2/Dm88, and Invader1 transposon families were upregulated over tenfold relative to controls (Figure 1). However, expression of all other transposons families was suppressed, and reduced expression persisted for 24hr (Figure 1A). In addition, the three elements that were initially over-expressed are progressively silenced during recovery. By contrast, the steady-state expression of most protein-coding genes did not change after heat shock, with the exception of the heat shock protein genes (HSPs; Figure S1A). This lack of change in gene expression was somewhat unexpected, as heat shock is known to suppress gene transcription (Jamrich, et al. 1977). However, a significant fraction of the transcriptome is deposited in developing oocytes for use during embryogenesis, and stable maternal RNAs may mask repressed transcription. Nonetheless, these findings indicate that acute heat shock leads to prolonged suppression of most transposon families in *D. melanogaster* ovaries.

**Figure 1:**
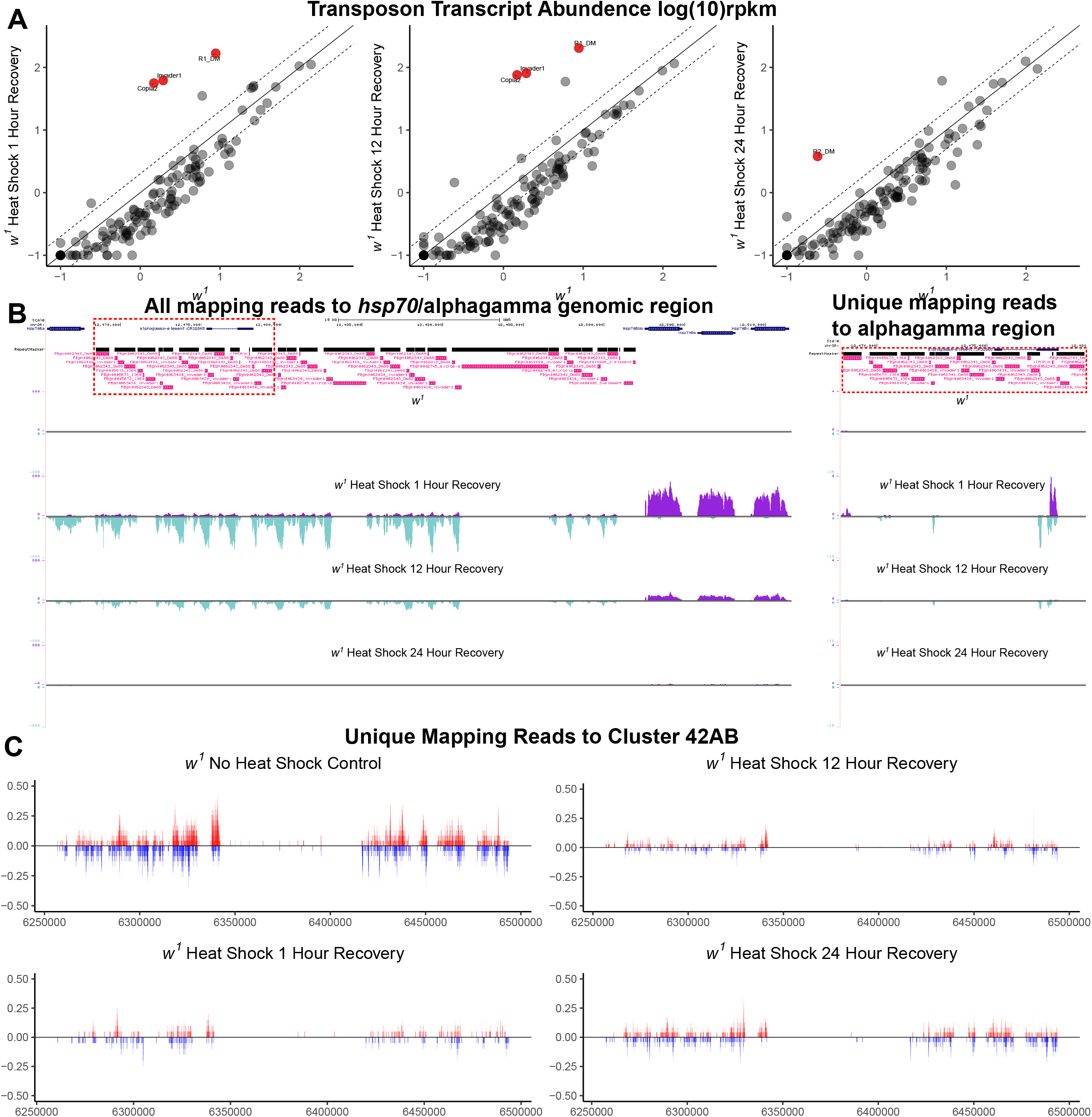
Acute heat shock downregulates all but a few transposons, and the major piRNA cluster 42AB. A. Example scatter plots comparing all mapping transposon family expression (rpkm, log10 scale) in *w^1^* control vs. samples heat shocked at 37°C for 1 hour and allowed to recover for 1, 12, and 24 hours. Each point represents a transposon family; families with an expression difference of 5-fold or greater are labeled and colored red, dotted line is 2-fold difference. B. A image from UCSC genome browser dm6 tracks for all mapping long RNA signal from the *hsp70B*/alpha gamma genomic region on chromosome 3R which contains numerous transposon insertions, including multiple invader1 and Dm88/Copia2 insertions. The tracks are in order: the reference sequence track (blue), repeat masker (black), transposon annotations (pink), *w^1^* no heat shock controls, *w^1^* heat shocked for 1 hour at 37°C allowed to recover for 1 hour, 12 hours, and 24 hours. Heat shock induces expression of *hsp70B* and Invader and Dm88/Copia2, which decrease expression coordinately. As all mapping sequences do not correspond to one location, we also show UCSC genome browser tracks from unique mapping long signal in the smaller insert from the region directly around the alpha gamma element. Tracks are in the same order, with different scales from above. Heat shock induces unique RNA expression from the alpha gamma element and the surrounding region. This unique expression indicates that at least some Invader1 and Dm88/Copia2 expression originates from this genomic region after heat shock. C. Unique mapping long RNA signal tracks for cluster 42AB in *w^1^* control, *w^1^* heat shocked for 1 hour at 37°C allowed to recover for 1 hour, 12 hours, and 24 hours. Heat shocked samples show lower expression across the cluster; this expression increases with time but does not recover to wild-type levels by 24 hours.

However, a small number of transposon families are transiently over-expressed in heat shocked ovaries. Two of these families, Copia2 and Invader1, are present in multiple copies within the 42AB piRNA cluster (Figure S1B). To determine if increased expression of these elements is associated with reduced piRNA levels, we sequenced small RNAs one hour after the shift to 37°C. At this point, R1, Copia2, and Invader 1 have reached maximal expression, and the remaining transposon families are silenced. However, there is no significant change in piRNAs mapping to the suppressed or over-expressed transposon families, or in piRNAs mapping to clusters (Figure S1C). The alpha gamma element lies between divergently transcribed *hsp70* genes and contains multiple copies of these transposons (Maside, et al. 2002; Kuan et al. 2009; Brewer-Jensen et al. 2016). Tellingly, expression of these families is highly correlated with expression of flanking *hsp70* genes, and with expression of unique sequences in the alpha gamma element (Figure 1B). We therefore speculate that activation of these transposon families is driven by heat shock induced transcription, not loss of piRNA silencing.

Heat shock significantly reduces expression of sequences that map uniquely to both strands of 42AB (Figure 1C), expression from both strands is partially restored at 24hr post heat shock, and this pattern of loss and recovery extends to most of the piRNA clusters (Figure S1D). The HP1 homolog Rhino anchors a complex that promotes transcription from both genomic strands (Klattenhoff et al. 2009; Volpe et al. 2001). To determine if heat shock disrupts Rhino binding, we used immunofluorescence and FISH to assay Rhino localization to the 42AB cluster. For this analysis, Rhino was immunolocalized, and the cluster was localized using a FISH probe to unique sequences immediately adjacent to 42AB. In control nurse cell nuclei, Rhino localizes to multiple foci, consistent with ChIP-seq and Cut and Run analyses of the genome wide distribution Rhino, and prominent Rhino foci are consistently found adjacent to the FISH signal marking one end of the 42AB clusters (Figure 2A). Immediately following heat shock, all Rhino foci are lost, including the major foci adjacent to 42AB (Figure 2B). 24 hours after heat shock, Rhino foci are restored, including foci adjacent to 42AB (Figure 2C). Heat shock thus triggers reversible loss of Rhino from germline clusters.

**Figure 2:**
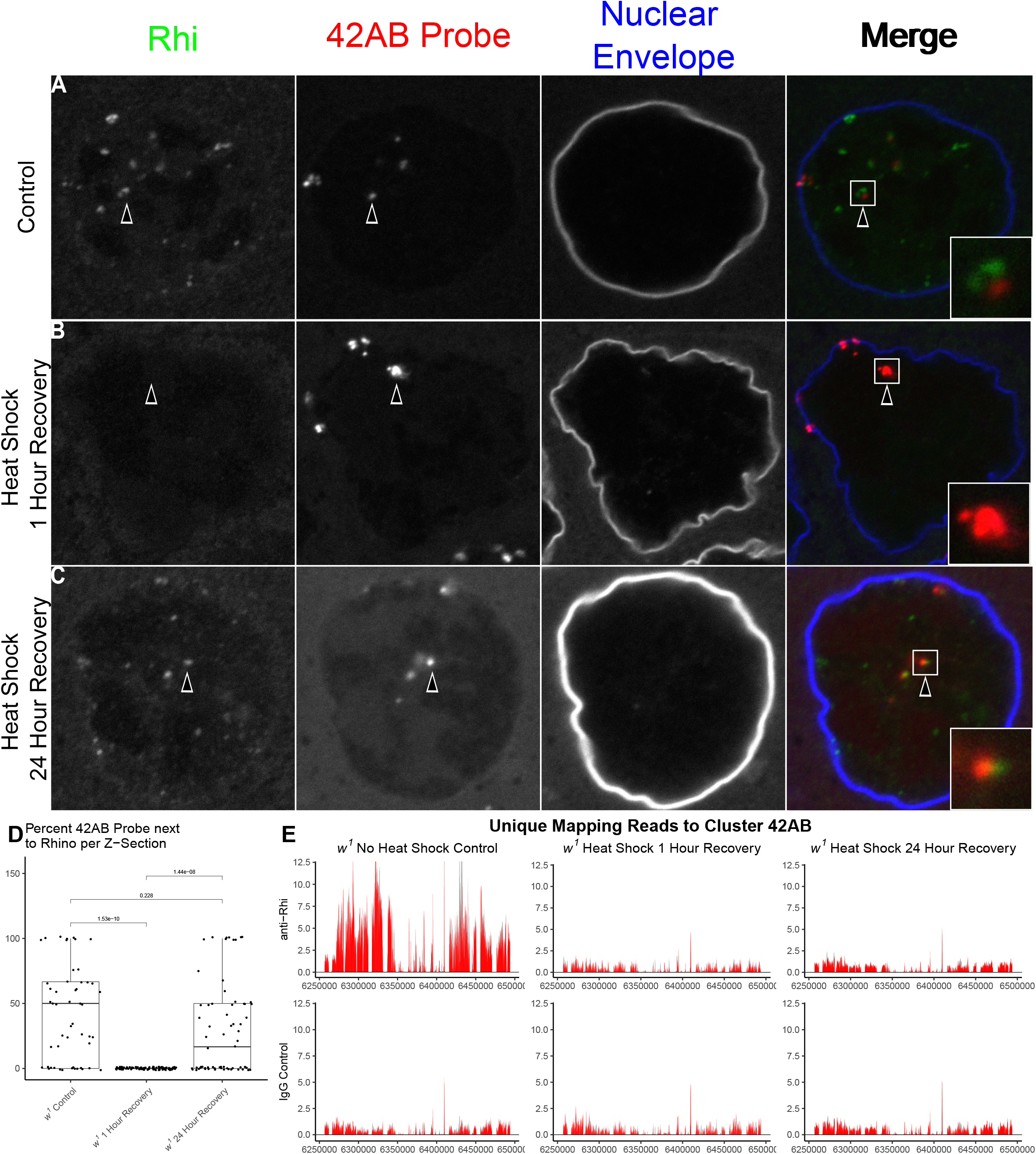
Heat shock causes the loss of Rhino localization. Rhino partially recovers after 24 hours. A-C. Confocal images of nurse cell nuclei in *w^1^* control (A) vs. *w^1^* heat shocked at 37°C for 1 hour and allowed to recover for 1 (B) and 24 (C) hours labeled for Rhino (anti-Rhino, green), DNA fish for unique sequence directly adjacent to cluster 42AB (red) and nuclear envelope (blue). Hollow arrows show foci in the merge inset. Rhino foci are next to 42AB adjacent foci before heat shock and 24 hours after, whereas Rhino foci are lost following heat shock and 1 hour recovery. D. Quantification of the relationship between Rhino and 42AB adjacent probe as a percentage of 42AB adjacent foci next to a Rhino foci per z-section (see methods for how next to was defined). Foci are defined as five standard deviations above the background. Each dot represents a single z-section. 100% indicates all 42AB adjacent foci were associated with Rhino foci, and 0% indicates no Rhino foci were next to 42AB adjacent foci. P-values for the indicated comparisons are above the bars. E. Cut and Run signal tracks for Rhino and IgG control uniquely mapping to 42AB in *w^1^* control showing Rhino and IgG signal at 42AB. Flies heat shocked at 37°C for 1 hour and allowed to recover for 1 and 24 hours showed no Rhino enrichment at 42AB vs. IgG control.

To quantify these observations, we used laser scanning confocal microscopy to optically section nurse cell nuclei and computationally defined 42AB and Rhino location by FISH and immunofluorescent foci, using a signal threshold of five standard deviations above the average for the field. Rhino foci “adjacent” to 42AB/FISH foci were identified based on signal overlap (see methods). This analysis confirmed that Rhino localization to 42AB recovers by 24hr after heat shock (Figure 2D).

To extend these cytological observations, we used Cut and Run (Ahmad 2018; Skene and Henikoff 2017) to determine the genome-wide distribution of Rhino before and after heat shock. Consistent with earlier observations, prior to heat shock, Rhino is enriched at 42AB (Figure 2E). Immediately after heat shock, by contrast, Rhino binding is undetectable across 42AB (Figure 2E). Surprisingly, Rhino binding remained undetectable 24 hours after heat shock, when transcription had partially recovered and Rhino cytologically localizes to 42AB (Figure 2E). piRNA cluster transcription collapses in *rhino* null mutants (Mohn et al. 2014), and partial restoration of cluster transcription from both strands after heat shock thus implies that functional complexes assemble during recovery from heat shock. Our inability to detect binding by Cut and Run, however, suggests that the nature of Rhino binding has changed, but the mechanistic basis for this change is unclear.

To further characterize heat shock control of cluster chromatin, we incubated female flies expressing GFP-Rhino at temperatures between 25°C and 37°C, dissected the ovaries, and imaged GFP in living egg chambers by laser scanning confocal microscopy. Rhino localizes to distinct nuclear foci following 1 hour incubation at temperatures up to 34°C, but foci were not detectable following a 1 hour incubation at 35°C (Figure S2A). To determine if Rhino localization is sensitive to absolute temperature or responds to a change in temperature, we raised GFP-Rhino flies at 18°C and shifted them to 30°C for 1 hour. Rhino foci remained following this treatment (Figure S2B). Assembly of cluster chromatin thus appears to sensitive to absolute temperature, not a temperature shift.

### Disassembly and reassembly of the piRNA precursor transcription and export machinery

Rhino anchors the Rhino-Deadlock-Cuff complex (RDC), which promotes non-canonical transcription from both strands and suppresses splicing and polyadenylation, generating precursor transcripts that are bound by the TREX complex and exported to the cytoplasm (Mohn et al. 2014; Zhang et al. 2018; Zhang, Wang, et al. 2014). Mutations in any single RDC component disrupt steady-state localization of the remaining proteins, but heat shock provides an opportunity to assay the kinetics of RDC disassembly. Therefore, we shifted female flies expressing GFP-Cuff to 37°C for 15min, 23mins, and 30mins, fixed the ovaries, immunolocalized Rhino, and quantified Cuff and Rhino signals in nuclear foci. As anticipated, Rhino and Cuff colocalize prior to heat shock (Figure 3A), and the signal for both proteins is highly correlated (scatter plot, Figure 3J). After a 15-minute heat shock, by contrast, Rhino is preferentially lost from nuclear foci, reflected in a shift to the left in the scatter plot comparing Cuff to Rhino signal (Figure 3J). Between 23min and 30min, Rhino is almost undetectable, and the Cuff signal has declined, but is still detected (Figure 3 B, C, J). We also find that 30 min heat shock displaces Deadlock from nuclear foci (Figure S3A). Heat shock thus temporally uncouples Cuff from Rhino localization.

**Figure 3:**
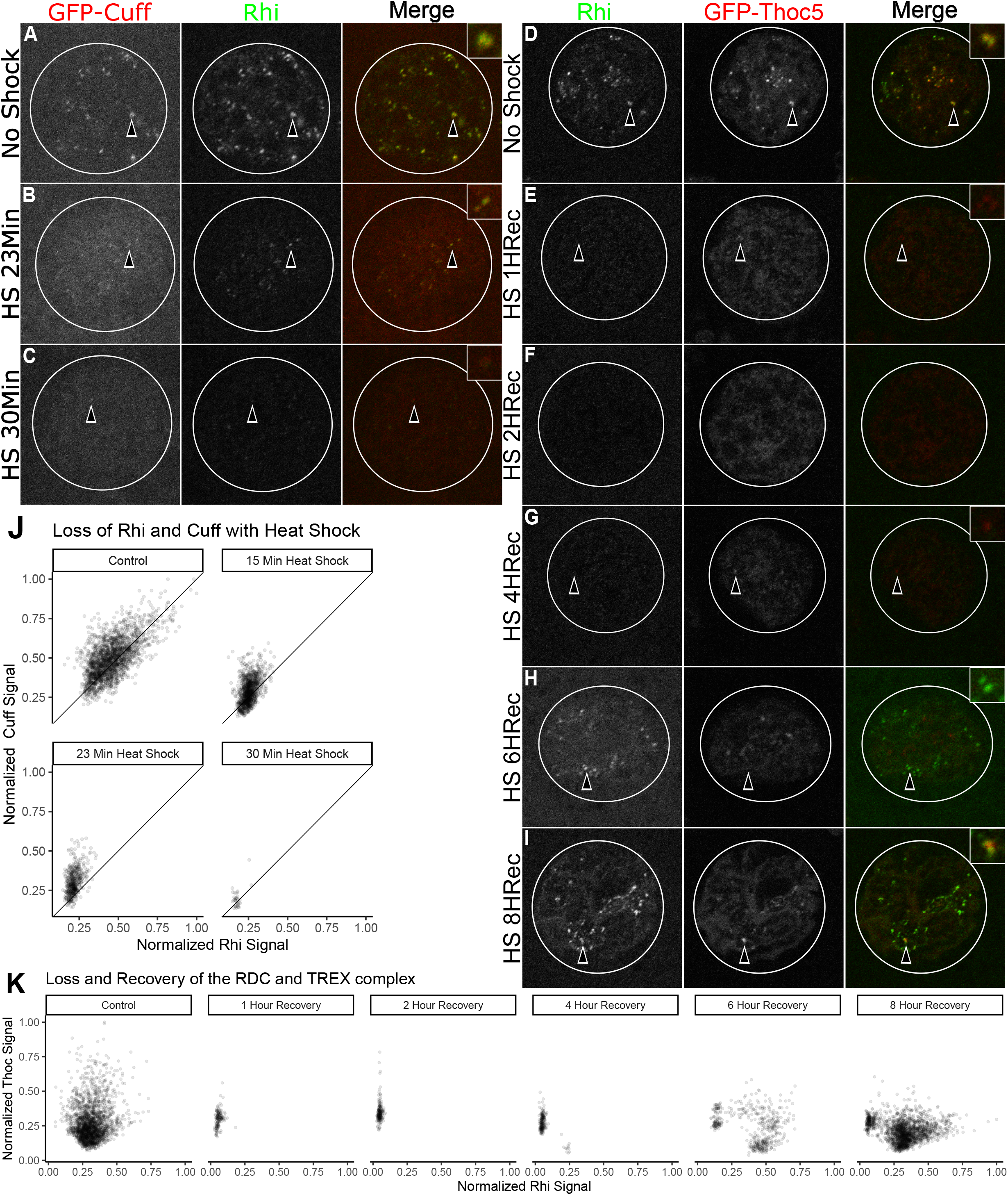
Disassembly and reassembly of the piRNA precursor transcription and export machinery. A. Confocal images of a nurse cell nucleus in GFP-Cuff control labeled for GFP-Cuff (anti-GFP, Red) Rhino (anti-Rhino, Green) and merge with arrows pointing at the inset which shows colocalized GFP-Cuff and Rhino. Levels were adjusted to achieve the maximum dynamic range. B. Same as above in GFP-Cuff heat shocked at 37°C for 23 minutes showing the reduction in Cuff and Rhino signal Same as above in GFP-Cuff heat shocked at 37°C for 30 minutes showing the loss of Cuff and Rhino signal. C. Confocal images of a nurse cell nucleus in GFP-Thoc5 control labeled for GFP-Thoc5 (anti-GFP, Red) Rhino (anti-Rhino, Green) and merge with arrows pointing at the inset shows colocalized GFP-Thoc5 and Rhino signal. Levels used the maximum signal as quantified below as the maximum value. D. Same as above in GFP-Thoc5 heat shocked at 37°C for 1 hour and allowed to recover for 1 hour showing loss of Rhino and Thoc5-GFP E. Same as above in GFP-Thoc5 heat shocked at 37°C for 1 hour and allowed to recover for 2 hours showing no recovery of Rhino and Thoc5-GFP F. Same as above in GFP-Thoc5 heat shocked at 37°C for 1 hour and allowed to recover for 4 hours showing no recovery of Rhino and Thoc5-GFP G. Same as above in GFP-Thoc5 heat shocked at 37°C for 1 hour and allowed to recover for 6 hours showing recovery of Rhino and not Thoc5-GFP. H. Same as above in GFP-Thoc5 heat shocked at 37°C for 1 hour and allowed to recover for 8 hours showing recovery of both Rhino and Thoc5-GFP. I. Scatterplot comparing the normalized signal of Rhino and GPF-Cuff foci in control with a 37°C heat shock for the indicated time. A single nurse cell nucleus was selected by DAPI staining to be the only nucleus in the optical section and then imaged in 1-micrometer steps. A minimum of four nuclei was used for each sample. Each point represents a focus of Rhino or GFP-Cuff five standard deviations above the background with the signal normalized to the maximum fluorescence in control. J. Scatterplot comparing the normalized signal of Rhino and GPF-Tho5 foci in control and after a 37°C heat shock for 1 hour and allowed to recover for the indicated time. The method used was the same as above. Each point represents focus of Rhino and/or GFP-Thoc5.

TREX binds to cluster transcripts and colocalizes with the RDC (Zhang et al. 2018). Thoc5 is a conserved component of the THO complex, a core component of TREX (Rehwinkel et al. 2004). To determine the impact of heat shock on TREX localization, we incubated flies expressing Thoc5-GFP at 37°C for 1hr and assayed Rhino and Thoc5-GFP localization 1, 2, 4, 6, and 8 hours after shifting back to 25°C. Figure 3 D-I shows representative images before heat shock and during recovery. Quantification of Rhino and Thoc5 localization to nuclear foci is shown in the scatter plots in Figure 3K. In control egg chambers, Rhino and Thoc5 colocalize to a subset of foci, but Rhino or Thoc5 individually dominate other foci. Following heat shock and one hour of recovery, Rhino is lost from nuclear foci, but someThoc5-GFP foci persists. Thoc5-GFP foci are essentially gone by 4hr of recovery, but Rhino foci have begun to recover. At this time point, and at all subsequent time points, Cuff colocalizes with Rhino (Image and Scatter plot Figure S3B and C), consistent with co-assembly. Rhino and Thoc5 co-localization is re-established 6hr into recovery, and increases at 8hr post heat shock (Scatter plots, Figure 3H). Venus tagged UAP56 showed a similar recovery pattern (Figure S3D). Recovery from heat shock thus appears to comprise assembly of the RDC and initiation of cluster transcription, followed by TREX binding to nascent transcripts.

### Heat shock control of cytoplasmic piRNA machinery

The RDC drives production of piRNA precursors that undergo TREX-dependent export from the nucleus and ping-pong processing and phased biogenesis in perinuclear nuage and at the surface of mitochondria. Aub and Vasa localize to nuage and are required for ping-pong amplification (Brennecke et al. 2007; Malone et al. 2009; Xiol et al. 2014). To determine the impact of heat shock on the ping-pong machinery, we analyzed Vasa and Aub localization after acute heat shock, using GFP fusions to follow protein distribution in live cells. Before heat shock, GFP-Vasa concentrates in nuage, and GFP-Aub is concentrated in nuage and present at lower levels in the cytoplasm (Figure 4). After heat shock, GFP-Aub is reduced at nuage and associates with large aggregates in the cytoplasm, and GFP-Vasa localizes to germline nuclei and shows reduced localization to nuage (Figure 4A and C). Mutations in *rhi, thoc7* or *uap56* disrupt Vasa and Aub localization to nuage (Zhang et al. 2018; Zhang et al. 2012; Klattenhoff et al. 2009) but do not lead to Vasa localization to the nucleus or Aub localization to cytoplasmic aggregates. The changes in Vasa and Aub localization thus appear to be a direct result of heat shock, not heat shock disruption of the RDC and TREX.

**Figure 4:**
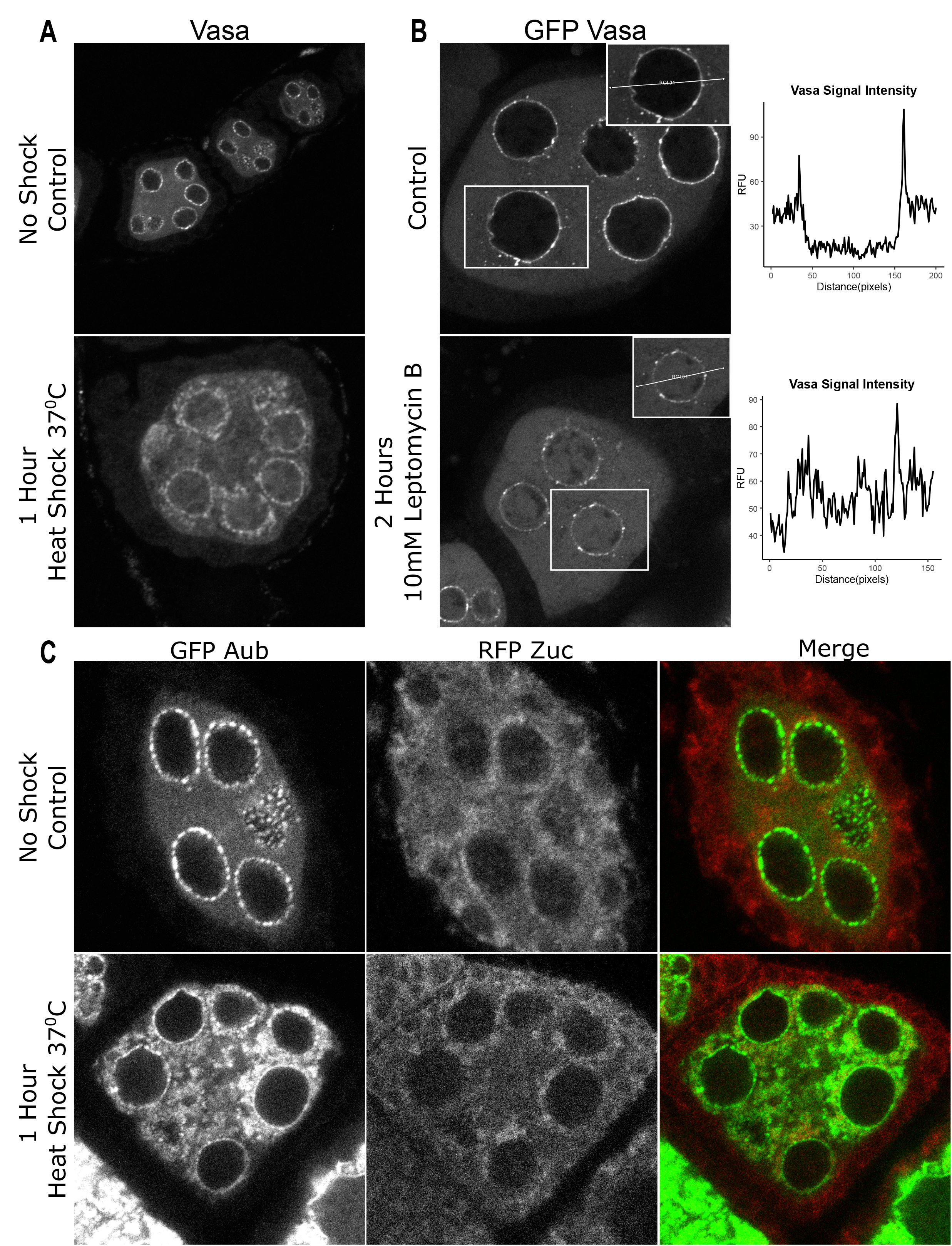
Acute heat shock causes accumulations of Vasa in the nucleus and Aub in mitochondria. A. Confocal images of an egg chamber stained for the piRNA pathway protein Vasa (anti-Vasa) in *w^1^* control and 1 hour after a 37°C heat shock for 1 hour showing Vasa staining inside the nucleus. B. Live confocal images of GFP-Vasa dissected egg chambers treated with 10mM leptomycin for 2 hours or DMSO control. Graphs to the right of images show line scans of GFP-Vasa signal from the single nurse cell nucleus shown in the inset. C. Confocal images of an egg chamber stained for the piRNA pathway proteins GFP-Aub (anti-GFP, green) and RFP-Zucchini (anti-RFP, red) in control and 1 hour after a 37°C heat shock for 1 hour showing Aub localization to perinuclear foci (nuage) and the additional cytoplasmic signal after heat shock while Zucchini signal is unchanged.

Vasa binds to cluster transcripts and localizes to nuage granules that are biased toward regions of the nuclear envelope opposite clusters (Zhang et al. 2012), Localization of Vasa to the nucleus on heat shock thus raised the intriguing possibility that Vasa cycles between the nucleus and nuage, perhaps facilitating precursor delivery to nuage. Cluster transcripts are exported from the nucleus through a CRM1-dependent process (ElMaghraby et al. 2019; Kneuss et al. 2019; Mendel and Pillai 2019). We therefore treated live egg chambers expressing GFP-Vasa with the CRM-1 inhibitor Leptomycin B and assayed localization (Prasad et al. 2007; Findley et al. 2003). After 2 hours of incubation in 10 µM Leptomycin B, GFP-Vasa localized to the nucleus and nuage (Figure 4B). The line scans in Figure 4B show Vasa signal across germline nuclei. In control egg chambers, the signal drops to near background levels in the nucleus, but the signal in the nucleus is comparable to the cytoplasmic signal following leptomycin B treatment (Figure 4B). Vasa thus appears to transiently cycle through the nucleus, which could facilitate cluster transcript export.

To further characterize the impact of heat shock on the piRNA biogenesis machinery, we assayed the localization of Ago3, Maelstrom, Squash, Armi, and Zuc. Ago3 localizes to nuage and functions with Aub and Vasa in ping-pong amplification (Wang et al. 2015; Webster et al. 2015; Malone et al. 2009). Maelstrom is concentrated in nuage and also found dispersed in the nucleus and cytoplasm, and suppresses canonical cluster transcription (Chang et al. 2019). Squash localizes to nuage and appears to function downstream of piRNA biogenesis in transposon silencing (Pane, et al. 2007; Malone et al. 2009). Armi localizes to nuage and mitochondria and functions with the mitochondrial nuclease Zuc in phased piRNA biogenesis (Ge et al. 2019). Using a combination of live imaging and immunofluorescence, we find that heat shock displaces Squash from nuage and leads to Maelstrom localization to cytoplasmic aggregates (Figure S4 A and B). However, heat shock has little impact on Ago3, Zuc, or Armi localization (Figure 4C and Figure S4 C, D). Heat shock thus disrupts piRNA cluster chromatin, the associated nuclear export machinery, and nuage localization of critical ping-pong amplification and silencing factors. By contrast, heat shock has relatively little impact on the localization of the phased piRNA biogenesis machinery.

### Genetic control of cluster chromatin assembly

piRNAs mapping to clusters are maternally deposited in the oocyte, and transient depletion of Piwi during embryogenesis has been reported to compromise piRNA cluster heterochromatin assembly in adults. By contrast, zygotic *piwi* mutations, which deplete this nuclear PIWI clade Argonaute through later stages of development, do not block cluster propagation or piRNA production in adults (Akkouche et al. 2017). These findings suggest that maternal piRNAs bound to Piwi direct cluster heterochromatin assembly in the embryo, and that clusters are then maintained by a Piwi-independent epigenetic mechanism. To determine the role of Piwi in cluster chromatin recovery following heat shock, we incubated *piwi^2/NLS^* mutant females at 37°C and assayed Rhino localization following a shift back to 25°C. The *piwi^NLS^* allele produces a protein lacking a nuclear localization signal (NLS), and is thus incapable of targeting chromatin (Klenov et al. 2011). Surprisingly, Rhino foci were present at 2hr post-heat shock and appeared to have fully recovered by 4hr post-heat shock (Figure 5A). In *w^1^* controls, by contrast, Rhino foci are just beginning to emerge after 4hr of recovery (Figure 5B). To quantify recovery, we determined the number of Rhino foci with signal five standard deviations above average in multiple nuclei at each time point. In *w^1^,* the first computationally-defined foci are observed at 4hr post-heat shock, and numbers have not fully recovered at 8hr. In *piwi^2/NLS^* mutants, numerous Rhino foci are detectable at 1hr post-heat shock, and localization has fully recovered by 4hrs. (Figure 5D). The *piwi* mutations thus *accelerate* recovery of Rhino localization.

**Figure 5:**
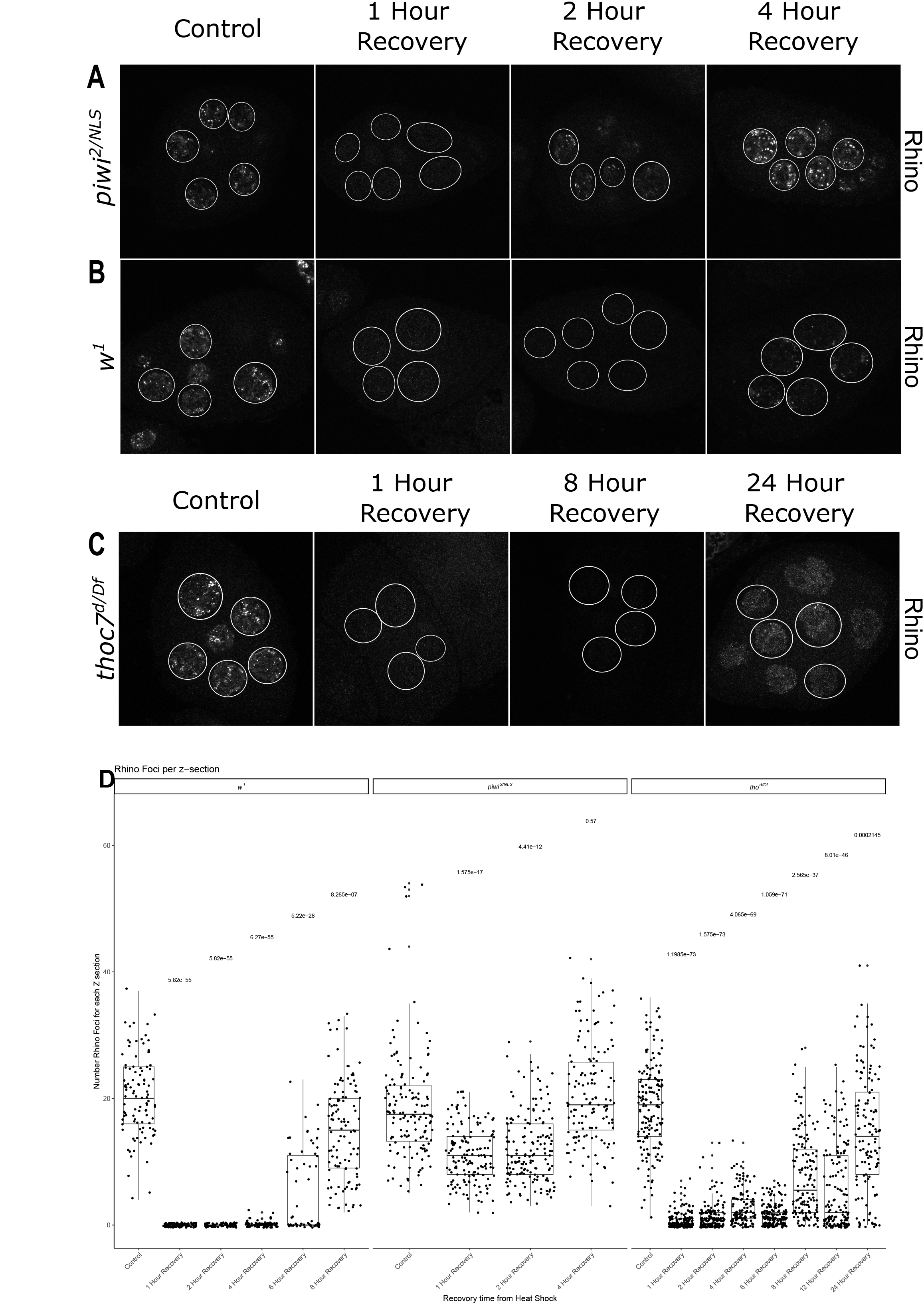
*piwi^2/NLS^* mutants accelerate Rhino foci recovery, and *thoc7^d/Df^* mutants delay recovery. A and B. Confocal images of egg chambers in *piwi^2/NLS^*(A) and *w^1^*(B) flies before heat shock, 1, 2, and 4 hours after a 1 hour 37°C heat shock stained for Rhino. Rhino foci are seen 4 hours after heat shock in *w^1^,* while *piwi^2/NLS^* show Rhino foci 2 hours after heat shock and numerous foci at 4 hours post heat shock. For clarity, circles show outlines of nurse cell nuclei in the field of view. C. Confocal images of egg chambers stained for Rhino in *thoc7^d/Df^* mutants before heat shock, 1, 8, and 24 hours after a 1 hour 37°C heat shock. For clarity, circles show outlines of nurse cell nuclei in the field of view. Eight hours after heat shock, many nuclei show no Rhino foci. Even 24 hours after heat shock, some nuclei show few Rhino foci. D. Quantification of the number of Rhino foci in *w^1^*, *piwi^2/NLS,^* and *thoc7^d/Df^* mutants after a 37°C heat shock for 1 hour and the indicated recovery time. Each dot represents the number of Rhino foci five standard deviations above background found in a single z section. A single nurse cell nucleus was selected by DAPI staining to be the only nucleus in the optical section and then imaged in 1-micrometer steps. A minimum of six nuclei was used for each sample. P-values compared to the no shock control are recorded above each bar.

piRNAs bound to Piwi function with Panoramix in piRNA-guided transcriptional silencing of transposons (Y. Yu et al. 2015; Sienski et al. 2015). To determine if the enhanced recovery of Rhino localization in *piwi* mutants is linked to transposon activation, we assayed recovery from heat shock in *pan^M/dD^* mutant ovaries. In this genetic background, Rhino recovery was comparable to controls (Figure S5A). These findings suggest that wild-type Piwi delays the reassembly of cluster chromatin following heat shock, through a process independent of transposon silencing.

The TREX complex localizes to clusters, binds cluster transcripts, and has been implicated in suppressing ectopic cluster chromatin assembly (Zhang et al. 2018). To determine the impact of TREX on cluster chromatin assembly, we assayed recovery of Rhino foci after heat shock in *thoc7^d/Df^* and *uap56^sz/28^* mutants. The *thoc7* gene encodes a component of the heteropentameric THO complex, which interacts with UAP56 in the TREX complex. However, THO and UAP56 also have TREX-independent functions (Hur and Chung 2016; Rehwinkel et al. 2004). In *thoc7^d/Df^* mutants, some Rhino foci are present 1hr after heat shock, but the number of foci did not begin to increase until 8hr of recovery, and wild-type numbers were not restored until 24hr after heat shock (Figure 5C). The *thoc7^d/Df^* mutation thus significantly delays Rhino recovery. By contrast, recovery of Rhino foci is comparable to *w^1^* in *uap56^sz/28^* mutants (Figure S5B). These findings suggest that the THO complex has a TREX-independent role in promoting Rhino localization.

### Maternal piRNAs are not required to license piRNA cluster chromatin

*piwi* mutations accelerate Rhino re-localization after heat shock, suggesting that Piwi and piRNAs are not required for cluster chromatin assembly in adult ovaries. After heat shock, however, we cannot detect Rhino binding by Cut and Run, indicating that wild-type cluster chromatin has not fully assembled. We therefore sought to independently test the role of piRNAs in cluster chromatin assembly.

Recently, Gebert et al. (2021) reported that deletion of the dominant germline piRNA cluster at 42AB does not disrupt transposon silencing or female fertility, despite removing approximately 30% of all piRNAs. Females homozygous for this deletion thus produce viable embryos that lack maternal piRNAs uniquely mapping to 42AB. We used this deletion to directly determine the role of maternal piRNAs in cluster chromatin assembly and piRNA biogenesis. For this analysis, we generated embryos lacking maternal piRNAs mapping to 42AB and carrying a wild-type copy of the cluster by crossing homozygous 42AB Df females to wild-type OreR males. The reciprocal cross generated genetically identical embryos with maternally deposited piRNAs (Figure 6A). We then assayed Rhino localization by Cut and Run and piRNA expression by small RNA-seq in the adult F1s.

**Figure 6:**
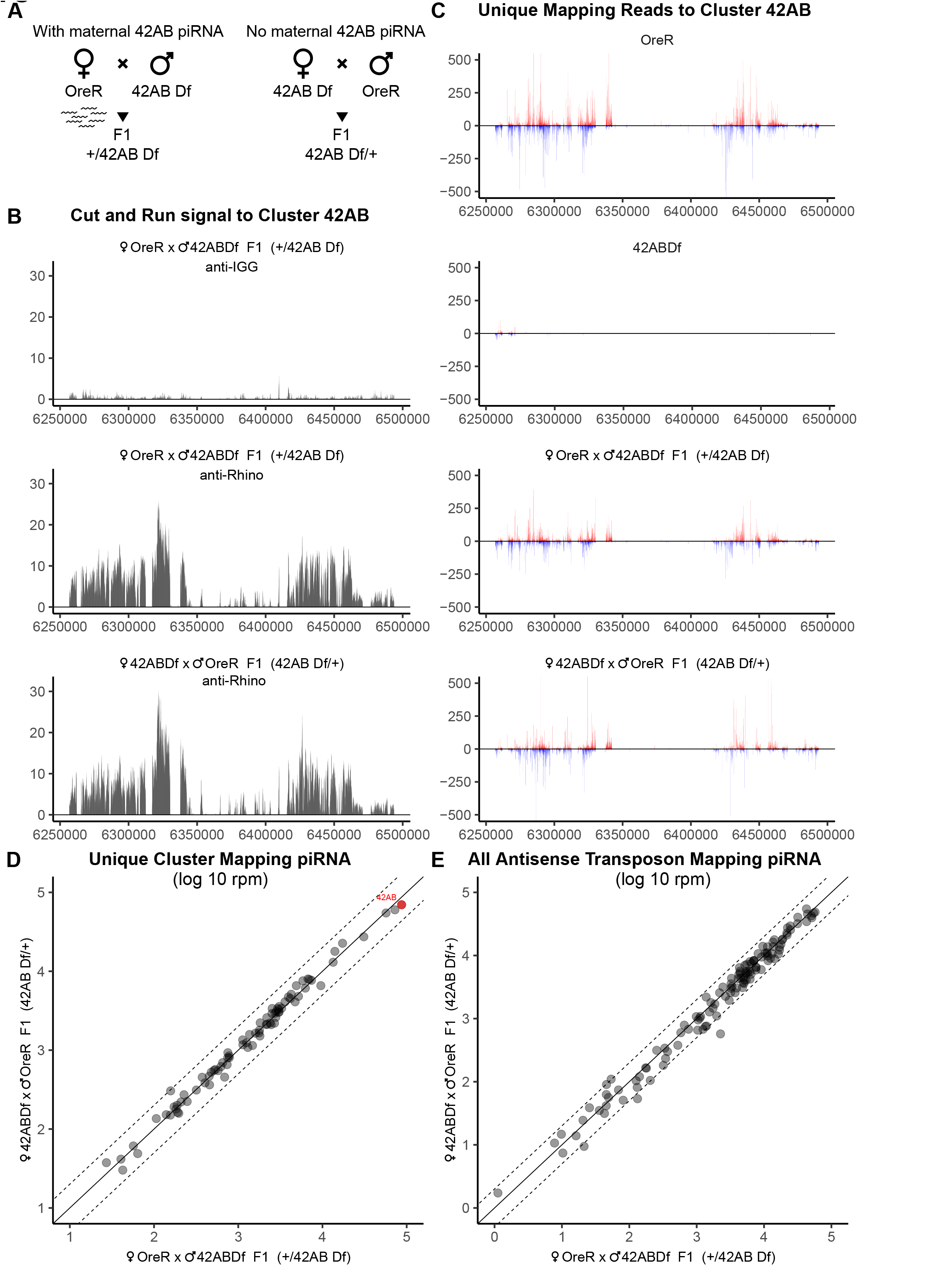
Maternal 42AB piRNAs are not required for licensing Rhino binding to and piRNA production from 42AB cluster. A. Experimental crosses: OreR females were crossed to 42AB Df males to produce F1 +/42AB Df heterozygotes that inherit maternal 42AB piRNAs. The reciprocal cross with 42AB Df females crossed to OreR males generates a genetically identical F1 population; however, they do not inherit maternal 42AB piRNAs. B. Cut and Run signal tracks at 42AB piRNA cluster for IgG control and anti-Rhino in F1 females with and without maternal piRNA. Graphs show similar enrichment of Rhino at 42AB piRNA cluster in both F1 populations. C. Uniquely mapping small RNA signal tracks against 42AB piRNA cluster for OreR, 42AB Df, and the two F1s with and without maternal 42AB piRNAs. No detectable differences between the two F1 populations were observed in three replicates. D. Example scatter plot comparing expression (rpkm log10) of mapping unique cluster piRNAs between two F1s show no global changes. Dotted line is 2 fold change. 42AB piRNA cluster is highlighted in red. E. Example scatter plot comparing expression (rpkm log10) of all transposon mapping piRNA between two F1s also show no global changes. Dotted line is 2 fold change.

Figure 6B and C show genome browser profiles of piRNA expression and Rhino binding across 42AB in the F1 progeny from these crosses. Remarkably, Rhino binding and piRNA production from 42AB is essentially identical in both. Quantification of cluster and transposon mapping piRNAs confirm these qualitative observations (Figure 6D and E). To confirm these findings, we performed a parallel cross using *w^1^* as the control strain. Again, piRNA production and Rhino binding were identical in the F1 progeny, independent of the maternal deposition of 42AB piRNAs (Figure S6A-E). We speculated that maternally-deposited piRNAs shared by 42AB and other clusters could license cluster chromatin. However, the alpha gamma genomic region consists mainly of transposable elements that are also imbedded in 42AB, and Rhino does not bind to this region, with and without maternal 42AB piRNAs (Figure S6F). These findings indicate maternal piRNAs are neither required nor sufficient to license piRNA cluster chromatin assembly.

## Discussion

Environmental stress leads to transposon activation in evolutionarily diverse systems and may generate genetic diversity that facilitates adaptive evolution. To drive evolution, mutations must be transmitted through the germline, and the piRNA pathway silences transposons during germline development. We, therefore, analyzed the impact of acute heat shock on the organization and function of the *Drosophila* piRNA pathway.

### Heat shock control of cluster chromatin

In *D. melanogaster,* the HP1 homolog Rhino binds to clusters and recruits the machinery that drives piRNA precursor production and TREX-dependent nuclear export. We show that heat shock leads to the rapid loss of Rhino, Deadlock, and Cuff localization to nuclear foci, followed by loss of the TREX complex (Figures 3 and S3). These cytological changes are associated with a decrease in cluster transcript expression from both strands of germline clusters, consistent with Rhino function in driving cluster transcription. Expression from clusters progressively increases during recovery from heat shock, which corresponds with the sequential restoration of Rhino and TREX localization to nuclear foci. Based on recovery of cluster transcription and dual FISH-immunofluorescence, Rhino localization of the 42AB cluster is restored (Figures 1, 2, 3). However, after 24hr of recovery, we do not detect Rhino binding to clusters, including 42AB, by Cut and Run, suggesting that wild type binding has not been re-established. Nonetheless, we find that acute heat shock has no impact on fertility, implying that cluster chromatin is functional.

### Heat shock control of the cytoplasmic piRNA biogenesis machinery

Following nuclear export, piRNA precursors undergo ping-pong and phased piRNA processing. The PIWI proteins Aub and Ago3, with the conserved DEAD box protein Vasa, are required for ping-pong amplification, and all three proteins localize to perinuclear nuage. Phased biogenesis requires Ago3, the conserved helicase Armi, and the nuclease Zuc. Ago3 localizes to nuage, Armi localizes to nuage and mitochondria, and Zuc associates with the mitochondrial membrane. Upon heat shock, Vasa rapidly localizes to the nucleus and is reduced at nuage, and Aub is displaced from nuage and localizes to granular material that lies adjacent to Zuc (Figure 4). Significantly, the CRM1 inhibitor leptomycin B also leads to the accumulation of Vasa in the nucleus, piRNA precursor export from the nucleus requires CRM1, and cluster transcripts co-precipitate with Vasa. These findings suggest that Vasa cycles between the nucleus and nuage and may directly facilitate export to piRNA precursors. The functional significance of Aub localization to structures near Zuc is unclear, as *aub* mutations do not disrupt phased biogenesis. However, Vasa is required for Aub localization to nuage, and reduced Aub localization to nuage on heat shock could be secondary to reduced Vasa localization.

### Two-phase response to heat shock?

Mutations in *rhi* and genes encoding the other RDC components disrupt cluster transcription and lead to a profound reduction in cluster mapping piRNAs, with broad increases in transposon expression. By contrast, loss of the RDC on heat shock reduces cluster transcript levels but does not reduce piRNAs mapping to clusters or transposons, and expression of most transposon families is suppressed. (Figure 1) Heat shock drives a broad loss of gene transcription (Jamrich, Greenleaf, and Bautz 1977; Lindquist 1986; Teves and Henikoff 2011). We, therefore, speculate that heat shock also suppresses transposon transcription, whereas piRNAs are stable and post-transcriptionally degrade the transposon transcripts that are present. The existing pool of piRNAs therefore may act as a buffer to allow time for the nuclear machinery to reassemble following acute heat shock. By contrast, prolonged heat stress has been reported to activate transposons and lead to transposition-induced mutations (Fanti et al. 2017; Cappucci et al. 2019). Prolonged disruption of the RDC with longer heat shock treatments may therefore allow time for turnover of relatively stable piRNAs, leading to transposon activation. A two-phase germline response to heat shock may function to maintain genome integrity in response to a relatively benign transient stress, but allow transposon mobilization and generation of genetic diversity on long term stress, facilitating adaptive evolution under conditions that compromise viability.

### Heat shock as a tool

The ability to reversibly disrupt cluster chromatin using heat shock provides a novel tool to probe the genetic control of cluster chromatin assembly. The sequential recovery of Rhino followed by TREX after heat shock is consistent with RDC function in promoting cluster transcription, producing precursors that are then bound by TREX and exported from the nucleus. How the RDC is directed to specific loci, by contrast, is not fully understood. piRNAs mapping to clusters and Piwi protein are maternally deposited in the oocyte, and transient depletion of Piwi during embryogenesis compromises, but does not block, Rhino localization and piRNA production in adults. By contrast, zygotic *piwi* mutations, which deplete the protein later in development, do not block Rhino localization or piRNA expression from clusters. Together, these observations suggest that maternal piRNAs bound to Piwi direct Rhino localization and cluster chromatin assembly during embryogenesis and that these heterochromatic domains are then propagated to the adult stage by a Piwi-independent epigenetic mechanism. However, we find that *piwi* mutations accelerate recovery of Rhino localization after heat shock. Conversely, *thoc7* mutations disrupt the TREX complex and delay recovery (Figure 5). Cluster assembly in the adult ovary thus appears independent of Piwi, and may be facilitated by TREX.

However, tight Rhino binding, as detected by Cut and Run, is not restored on recovery from heat shock, and assembly of structurally wild type cluster chromatin could require piRNAs bound to Piwi. To rigorously test this hypothesis, we used a deletion that removes the major germline cluster at 42AB. This deletion is viable and fertile, and embryos derived from homozygous females thus lack maternally deposited piRNA mapping specifically to this cluster. Remarkably, 42AB piRNA production and Rhino binding in the adult offspring of homozygous deletion females mated to wild type OreR or *w^1^* males are essentially identical to the offspring of the reciprocal cross. We speculated that piRNAs that map to 42AB and additional loci could initiate cluster chromatin assembly. However, Rhino does not localize in other genome regions matching these “multi-mapping” piRNAs, in wild type ovaries or ovaries from the offspring crosses with the deficiency. Cluster chromatin assembly is, therefore, independent of maternal piRNAs.

Mutations in *thoc7* delay the reassembly of cluster chromatin following heat shock, and the Thoc7 protein is a component of the RNA binding THO complex. Co-transcriptional binding of the THO to cluster transcripts, presumably generated by Rhino-independent canonical transcription, could therefore initiate cluster chromatin assembly. Promoter elements from transposon fragments within clusters could generate transcripts that are bound by the THO, but how the resulting transcripts are differentiated from gene transcripts is unclear. Reversible assembly of cluster chromatin following heat shock provides a novel tool to address these questions, and demonstrates that the piRNA pathway is dynamic and rapidly responds to the environment.

## Methods

### Resources Table

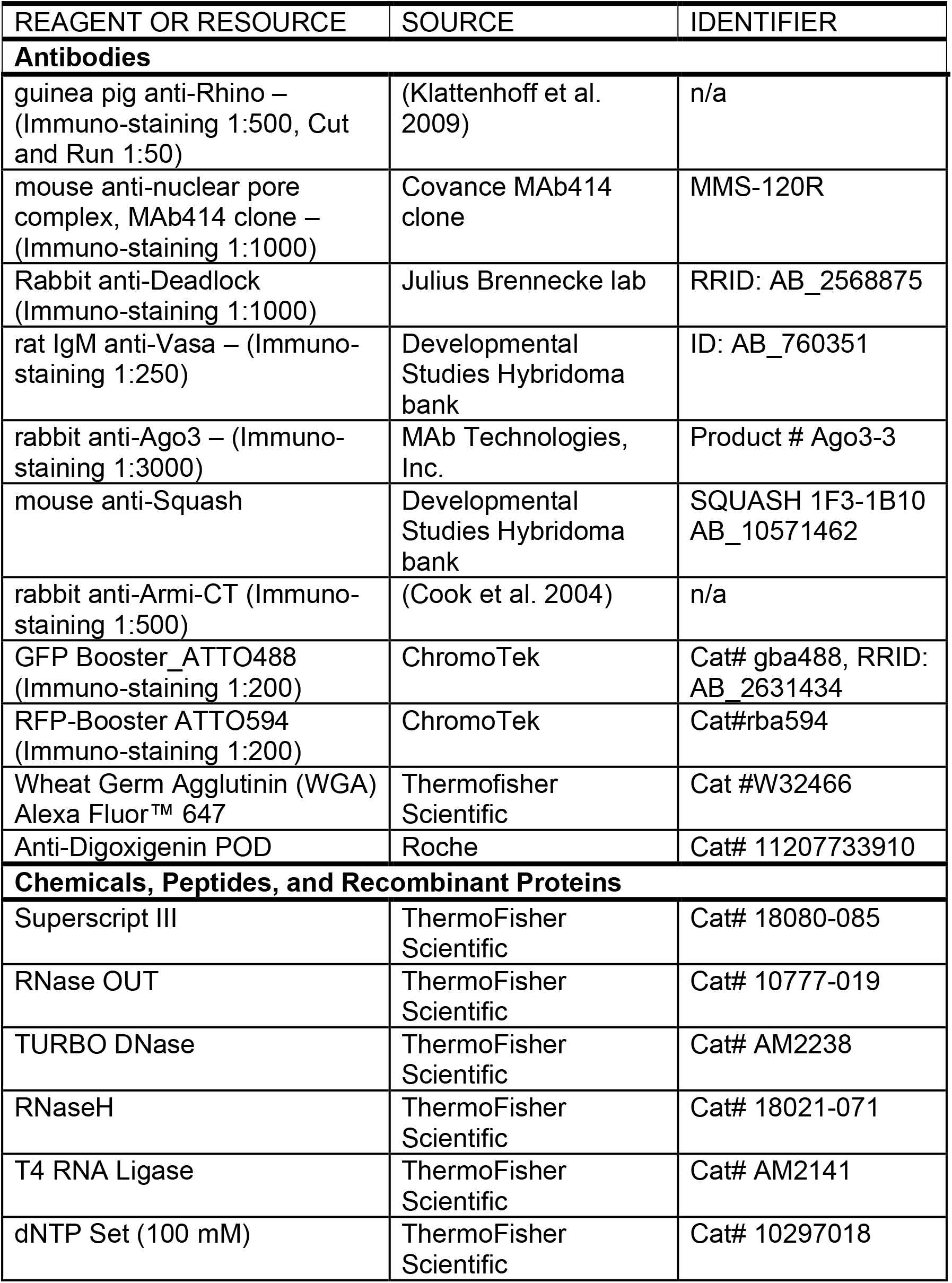
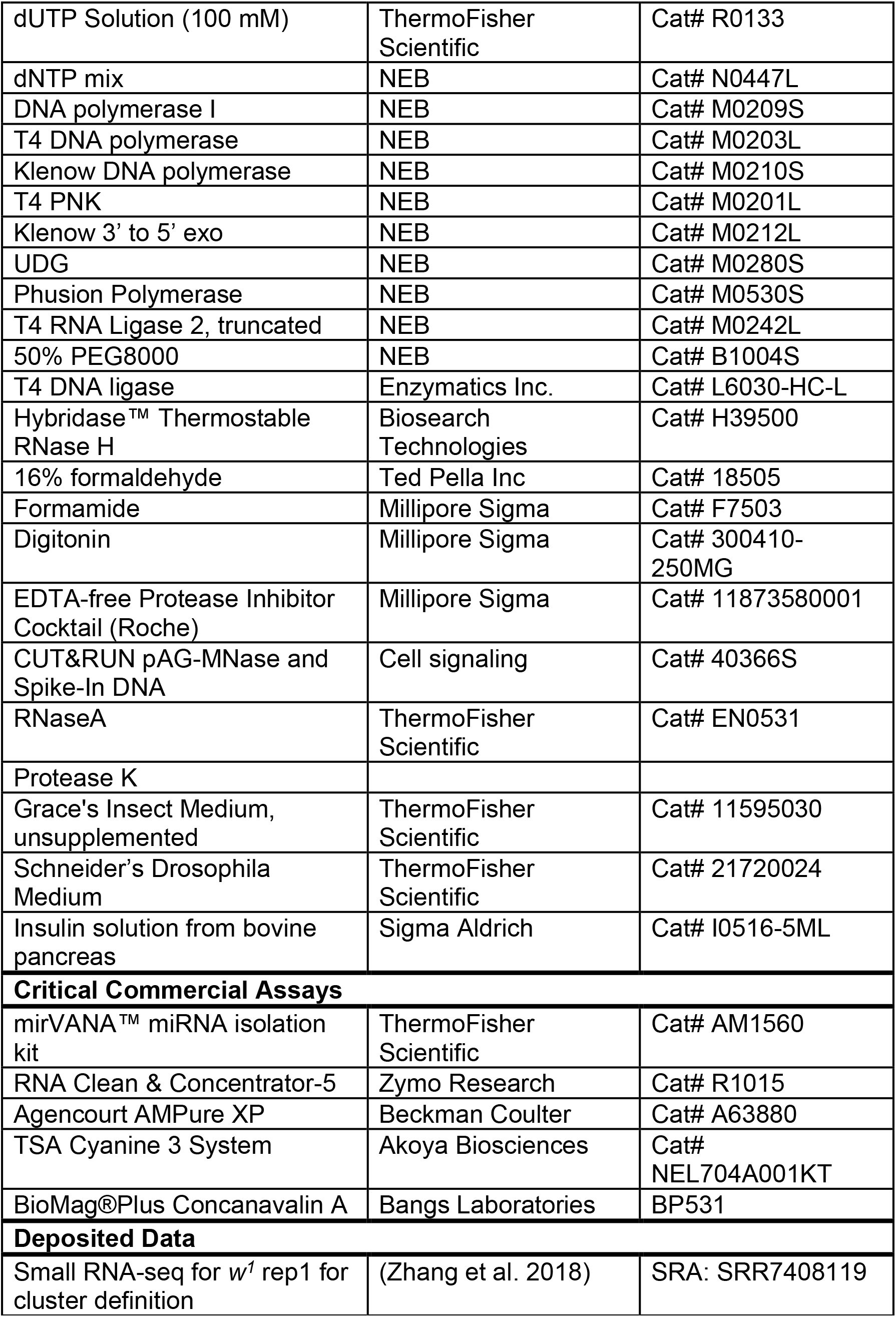
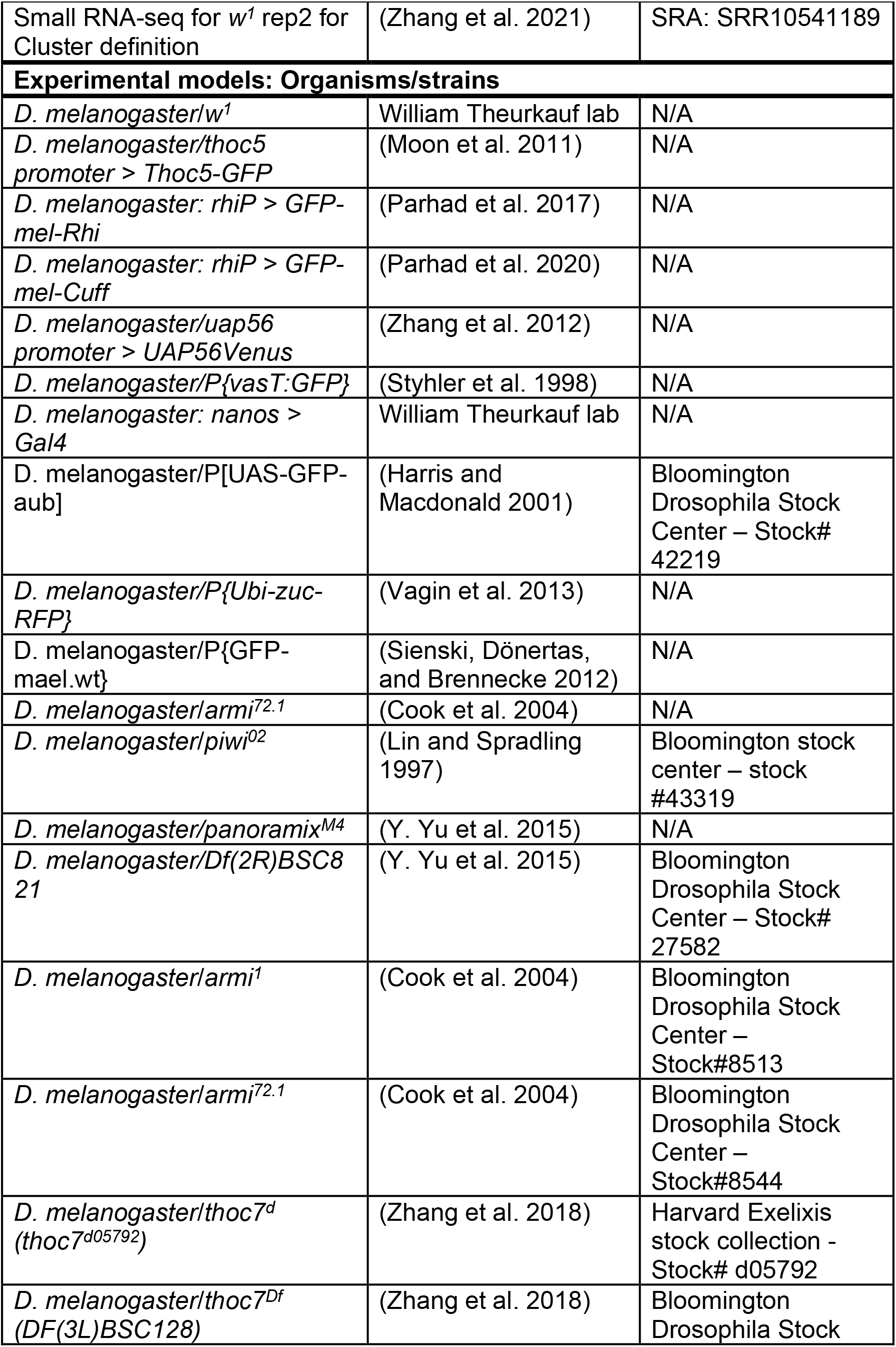
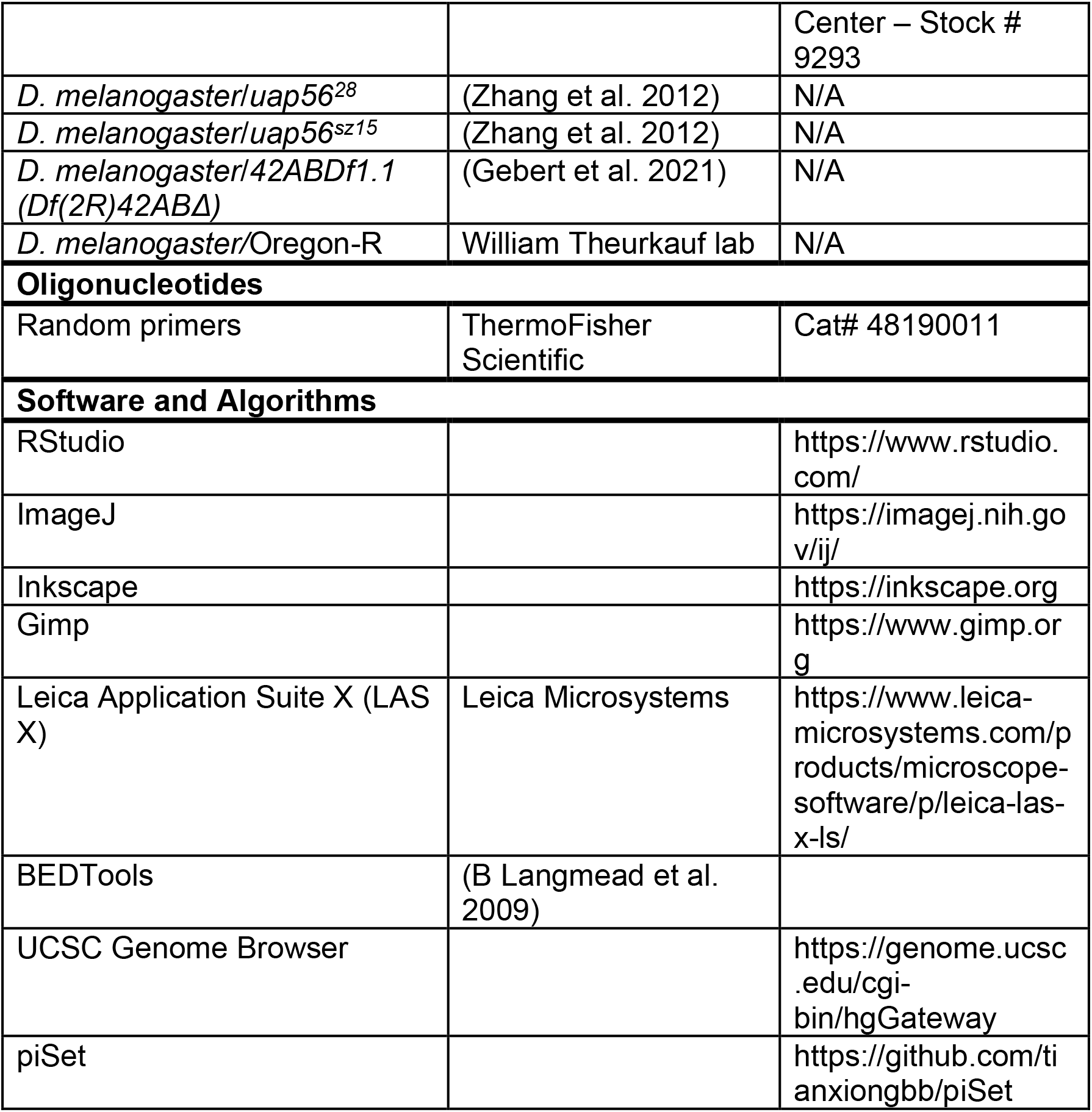

| REAGENT OR RESOURCE | SOURCE | IDENTIFIER |
|---|---|---|
| **Antibodies** |  |  |
| guinea pig anti-Rhino – (Immuno-staining 1:500, Cut and Run 1:50) | (Klattenhoff et al. 2009) | n/a |
| mouse anti-nuclear pore complex, MAb414 clone – (Immuno-staining 1:1000) | Covance MAb414 clone | MMS-120R |
| Rabbit anti-Deadlock (Immuno-staining 1:1000) | Julius Brennecke lab | RRID: AB_2568875 |
| rat IgM anti-Vasa – (Immuno-staining 1:250) | Developmental Studies Hybridoma bank | ID: AB_760351 |
| rabbit anti-Ago3 – (Immuno-staining 1:3000) | MAb Technologies, Inc. | Product # Ago3-3 |
| mouse anti-Squash | Developmental Studies Hybridoma bank | SQUASH 1F3-1B10<br>AB_10571462 |
| rabbit anti-Armi-CT (Immuno-staining 1:500) | (Cook et al. 2004) | n/a |
| GFP Booster_ATTO488 (Immuno-staining 1:200) | ChromoTek | Cat# gba488, RRID: AB_2631434 |
| RFP-Booster ATTO594 (Immuno-staining 1:200) | ChromoTek | Cat#rba594 |
| Wheat Germ Agglutinin (WGA) Alexa Fluor™ 647 | Thermofisher Scientific | Cat #W32466 |
| Anti-Digoxigenin POD | Roche | Cat# 11207733910 |
| **Chemicals, Peptides, and Recombinant Proteins** |  |  |
| Superscript III | ThermoFisher Scientific | Cat# 18080-085 |
| RNase OUT | ThermoFisher Scientific | Cat# 10777-019 |
| TURBO DNase | ThermoFisher Scientific | Cat# AM2238 |
| RNaseH | ThermoFisher Scientific | Cat# 18021-071 |
| T4 RNA Ligase | ThermoFisher Scientific | Cat# AM2141 |
| dNTP Set (100 mM) | ThermoFisher Scientific | Cat# 10297018 |
| dUTP Solution (100 mM) | ThermoFisher Scientific | Cat# R0133 |
| dNTP mix | NEB | Cat# N0447L |
| DNA polymerase I | NEB | Cat# M0209S |
| T4 DNA polymerase | NEB | Cat# M0203L |
| Klenow DNA polymerase | NEB | Cat# M0210S |
| T4 PNK | NEB | Cat# M0201L |
| Klenow 3' to 5' exo | NEB | Cat# M0212L |
| UDG | NEB | Cat# M0280S |
| Phusion Polymerase | NEB | Cat# M0530S |
| T4 RNA Ligase 2, truncated | NEB | Cat# M0242L |
| 50% PEG8000 | NEB | Cat# B1004S |
| T4 DNA ligase | Enzymatics Inc. | Cat# L6030-HC-L |
| Hybridase™ Thermostable RNase H | Biosearch Technologies | Cat# H39500 |
| 16% formaldehyde | Ted Pella Inc | Cat# 18505 |
| Formamide | Millipore Sigma | Cat# F7503 |
| Digitonin | Millipore Sigma | Cat# 300410-250MG |
| EDTA-free Protease Inhibitor Cocktail (Roche) | Millipore Sigma | Cat# 11873580001 |
| CUT&RUN pAG-MNase and Spike-In DNA | Cell signaling | Cat# 40366S |
| RNaseA | ThermoFisher Scientific | Cat# EN0531 |
| Protease K |  |  |
| Grace's Insect Medium, unsupplemented | ThermoFisher Scientific | Cat# 11595030 |
| Schneider's Drosophila Medium | ThermoFisher Scientific | Cat# 21720024 |
| Insulin solution from bovine pancreas | Sigma Aldrich | Cat# I0516-5ML |
| **Critical Commercial Assays** |  |  |
| mirVANA™ miRNA isolation kit | ThermoFisher Scientific | Cat# AM1560 |
| RNA Clean & Concentrator-5 | Zymo Research | Cat# R1015 |
| Agencourt AMPure XP | Beckman Coulter | Cat# A63880 |
| TSA Cyanine 3 System | Akoya Biosciences | Cat# NEL704A001KT |
| BioMag®Plus Concanavalin A | Bangs Laboratories | BP531 |
| **Deposited Data** |  |  |
| Small RNA-seq for *w<sup>1</sup>* rep1 for cluster definition | (Zhang et al. 2018) | SRA: SRR7408119 |
| Small RNA-seq for *w<sup>1</sup>* rep2 for Cluster definition | (Zhang et al. 2021) | SRA: SRR10541189 |
| **Experimental models: Organisms/strains** |  |  |
| *D. melanogaster/w<sup>1</sup>* | William Theurkauf lab | N/A |
| *D. melanogaster/thoc5 promoter > Thoc5-GFP* | (Moon et al. 2011) | N/A |
| *D. melanogaster: rhiP > GFP-mel-Rhi* | (Parhad et al. 2017) | N/A |
| *D. melanogaster: rhiP > GFP-mel-Cuff* | (Parhad et al. 2020) | N/A |
| *D. melanogaster/uap56 promoter > UAP56Venus* | (Zhang et al. 2012) | N/A |
| *D. melanogaster/P{vasT:GFP}* | (Styhler et al. 1998) | N/A |
| *D. melanogaster: nanos > Gal4* | William Theurkauf lab | N/A |
| D. melanogaster/P[UAS-GFP-aub] | (Harris and Macdonald 2001) | Bloomington Drosophila Stock Center – Stock# 42219 |
| *D. melanogaster/P{Ubi-zuc-RFP}* | (Vagin et al. 2013) | N/A |
| D. melanogaster/P{GFP-mael.wt} | (Sienski, Dönertas, and Brennecke 2012) | N/A |
| *D. melanogaster/armi<sup>72.1</sup>* | (Cook et al. 2004) | N/A |
| *D. melanogaster/piwi<sup>02</sup>* | (Lin and Spradling 1997) | Bloomington stock center – stock #43319 |
| *D. melanogaster/panoramix<sup>M4</sup>* | (Y. Yu et al. 2015) | N/A |
| *D. melanogaster/Df(2R)BSC821* | (Y. Yu et al. 2015) | Bloomington Drosophila Stock Center – Stock# 27582 |
| *D. melanogaster/armi<sup>1</sup>* | (Cook et al. 2004) | Bloomington Drosophila Stock Center – Stock#8513 |
| *D. melanogaster/armi<sup>72.1</sup>* | (Cook et al. 2004) | Bloomington Drosophila Stock Center – Stock#8544 |
| *D. melanogaster/thoc7<sup>d</sup> (thoc7<sup>d05792</sup>)* | (Zhang et al. 2018) | Harvard Exelixis stock collection - Stock# d05792 |
| *D. melanogaster/thoc7<sup>Df</sup> (DF(3L)BSC128)* | (Zhang et al. 2018) | Bloomington Drosophila Stock Center – Stock # 9293 |
| *D. melanogaster/uap56<sup>28</sup>* | (Zhang et al. 2012) | N/A |
| *D. melanogaster/uap56<sup>sz15</sup>* | (Zhang et al. 2012) | N/A |
| *D. melanogaster/42ABDf1.1 (Df(2R)42ABΔ)* | (Gebert et al. 2021) | N/A |
| *D. melanogaster/*Oregon-R | William Theurkauf lab | N/A |
| **Oligonucleotides** |  |  |
| Random primers | ThermoFisher Scientific | Cat# 48190011 |
| **Software and Algorithms** |  |  |
| RStudio |  | https://www.rstudio.com/ |
| ImageJ |  | https://imagej.nih.gov/ij/ |
| Inkscape |  | https://inkscape.org |
| Gimp |  | https://www.gimp.org |
| Leica Application Suite X (LAS X) | Leica Microsystems | https://www.leica-microsystems.com/products/microscope-software/p/leica-las-x-ls/ |
| BEDTools | (B Langmead et al. 2009) |  |
| UCSC Genome Browser |  | https://genome.ucsc.edu/cgi-bin/hgGateway |
| piSet |  | https://github.com/tianxiongbb/piSet |

### Fly strains and husbandry

Fly strains used are in the Key Resources table. All flies were maintained at 25°C on cornmeal medium. 1-2 day old females were collected and fed on yeast for 2 days. Heat shock was carried on 2-4 day old flies with yeast for 1 hour in a water bath kept at 37°C or at the stated temperature. Flies were allowed to recover for 1 hour after heat shock unless otherwise stated. Ovaries were collected from aged matched 2-4 day old females for a no heat shock control.

### Immunofluorescence

Staining of fix ovaries was performed as previously described with buffer A protocol in Theurkauf 1994 and Zhang et al. 2012. Briefly, 2-4 day old female ovaries were dissected in Robb’s media, fixed in 4% formaldehyde for 10 minutes, washed, incubated in primary overnight, washed, incubated in fluorophore conjugated secondary and DAPI for DNA staining overnight if required. If staining for nuclear envelope, incubated for 30 minutes at room temperature with wheat germ agglutinin with a final concentration of 5ng/ul, washed and mounted on the slide. Primary antibody concentrations used were as follows: guinea pig anti-Rhino 1:500, rat IgM anti-Vasa 1:500, rabbit anti-Ago3 1:3000, mouse anti-Squash 1:30, GFP and RFP booster at 1:200. See Key Resources Table for antibody details.

Live ovaries dissection was carried out using media described in Prasad et al. 2007 and were mounted directly on a coverslip in halocarbon oil for live imaging. Treatment with Leptomycin B was carried out in the media mix in Prasad et al. 2007 at 25°C for the indicated time in a rocking incubator.

### Immuno-DNA Fish

Immunofluorescence staining was performed the same as above up until completion of secondary conjugation overnight. After secondary incubation, samples were washed in BAT buffer for 10 mins, then fixed in Buffer A Fixative containing 4% formaldehyde for 10 mins as described in the immunofluorescence protocol. Samples were rinsed in 2xSSCT (2x SSC, 0.1% Tween-20) three times to exchange buffering system and subsequently washed at room temp for 10 mins with 2xSSCT+20% formamide, then 2xSSCT+40% formamide, and lastly 2xSSCT+50% formamide to gradually increase formamide concentration. Fresh 2xSSCT+50% formamide was added to the sample and incubated at 37°C water bath for 2 hours to allow for equilibration. Solution was removed and replaced with 0.3ug probe (the probe to 42AB was against a unique region directly adjacent to 42AB itself as 42AB is repetitive may bind else in the genome) and in buffer containing 1xSSC, 50% Formamide, and 10% (m/v) dextran sulfate for each sample, then denatured at 95°C for 5 mins, and allowed to hybridize overnight at 37°C. Samples were then washed for 20 mins with pre-warmed 2xSSCT+50% formamide at 37°C three times, followed by 10 mins washes at room temperature with solutions in the following order: 2xSSCT+40% formamide, 2xSSCT+20% formamide, 2xSSCT wash three times, PBST (PBS+0.2% tween-20) wash 2 times. anti-DIG POD conjugation (Roche) and Tyramide signal amplification (Akoya Biosciences) was performed according to manufacturer’s instruction. DAPI staining was performed simultaneously with Cy3 addition when required, followed by 3 washes with PBS-0.05% Triton X for 5 mins before mounting on slides. All images were taken a Lieca TCS SP8 confocal microscope and line scans were produced using Lieca Application Suite X.

### Automated Image Analysis

To quantify the presence of Rhino adjacent to 42AB foci, we developed an automated and unbiased image analysis pipeline using ImageJ. z-sections of nurse cell nuclei stained for Rhino and 42AB-DNA were taken using confocal microcopy. Signal was defined as 5 standard deviations above the mean signal for each marker. Number of 42AB foci were counted for each z section and designated as regions of interest using analyze particles. To measure number of 42AB adjacent to Rhino, we looked for overlap between Rhino signal in each region of interest. Rhino and 42AB signal are usually just shy of colocalization. To ensure that we capture adjacent localization of these two markers, we expanded the Rhino signal equally in all directions and measured overlap with each exaptation. A range of 0-6 pixel expansion were tested and all expansions showed the same trend. Expansion 3 was chosen for the main figure.

For Thoc7-Rhi and Cuff-Rhi colocalization analysis z-stacks were taken of nurse cell nuclei stained for two different nuclear markers. Images were analyzed in ImageJ, where signal was defined the same way as above. Signal for the two markers were merged and used to pick particles to define region of interest. For every focus identified, the signal intensity for each marker was measured.

To count Rhino foci z-stacks were acquired of nurse cell nucleus stained for Rhino and signal was defined the same as above. Rhino foci for each z section was defined using analyze particle and subsequently counted.

### Small RNA seq

Total RNA seq was performed as in Zhang et al. 2021, Briefly MirVana kit (Ambion) was used to extract total RNA from 2-4 day old female ovaries. 18-29 nucleotide small RNAs were isolated using polyacrylamide gel purification. Sequencing libraries were constructed as follows: 2S rRNA depletion, 3’ adaptor ligation, gel purification, 5’ adaptor ligation, gel purification, reverse transcription, and PCR amplification. Libraries were sequenced using Illumina Nextseq platform and sequencing analysis mapping to *dm6* was performed using piSet_srnaseq pipeline on GitHub.

### Long RNA seq

Total RNA sequencing was performed as described Zhang et al. 2021 Total RNA was extracted from 2-4 day old female ovaries using the MirVana kit (Ambion). rRNA was depleted using antisense rRNA oligo hybridization with subsequent RNase H digestion. Libraries were made as follows: RNA Fragmentation, Reverse Transcription, dUTP incorporation, end repair, size selection with Ampure XP beads, A-tailing, Adaptor ligation, UDG treatment and PCR amplification. Sequencing was performed using the Illumina Nextseq and sequencing analysis mapping to *dm6* was performed using piSet_rnaseq pipeline on GitHub.

### Cut and Run

Cut and Run protocol was adapted form Ahmad, 2018 (<u>dx.doi.org/10.17504/protocols.io.umfeu3n</u>) (Ahmad 2018; Skene and Henikoff 2017). A brief description and modification are as follows. Ovaries were dissected from 2-4 day old females in Robb’s medium, bound to ConA beads, incubated overnight in primary antibody, guinea pig anti-rhino at 1:50 dilution or guinea pig IgG, and incubated with pAMA. DNA was digested for 1 hour before reaction was stopped. DNA recovered from supernatant fraction was used to prepare sequencing libraries by end Repair, A-tailing, Adaptor ligation, and then PCR amplification. Libraries were sequenced using Illumina Nextseq. Sequencing analysis mapping to dm6 was performed using piSet_chipseq pipeline on GitHub.

### Bioinformatics analysis

The bioinformatic analysis was previously described in Zhang et al. (2021). The *Drosophila* reference genome (*dm6*), rRNA sequences, gene annotations, and hairpin sequences were obtained from Flybase (Version 6.13). Transposon consensus sequence were obtained from Repbase (Bao, Kojima, and Kohany 2015).

### piRNA cluster annotation

piRNA clusters were defined used a similar method as T. Yu et al. 2019. *w^1^* control small RNA-seq data from Zhang et al. (2021) was used to annotate piRNA clusters on *dm6* (See Key Resource table). Data mapping to rRNA, miRNA, tRNA, snRNA, and snoRNA were removed and the remaining 24–32 nt small RNA reads that mapped to the *dm6* genome were considered piRNAs. These piRNAs were then assigned to 20 kb sliding windows (with a 1 kb step). Windows containing more than 100 piRNAs per million uniquely mapped piRNAs were considered potential piRNA clusters. 20-kb genomic windows containing less than 200 distinct read or species were also filtered out to remove false positives resulting from un-annotated miRNA, rRNA, tRNA, snRNA, and snoRNA that mostly produce reads of the same sequence. Additionally, we calculated the first-nucleotide content for each 20-kb window and discarded those with 1U/10A percentage less than 50%. The remaining contiguous 20-kb windows were deemed putative piRNA clusters. Lastly, we performed manual curation for putative piRNA clusters using piRNA profile. piRNA clusters were classified as uni-strand and dual-strand piRNA clusters based on direction of the piRNAs produced.

### RNA-Seq Analysis

piSet_rnaseq pipeline from GitHub was used to analyze long RNA sequences reads. Raw reads were mapped to rRNA sequence using Bowtie2 (Version 2.2.5) with default settings (Ben Langmead and Salzberg 2012). Remaining reads were mapped to the *Drosophila* genome (*dm6*) and transposon consensus sequences using STAR (Version 2.5.2b) and Hisat2 with default parameters (Dobin et al. 2013; Kim, Langmead, and Salzberg 2015). The transcript abundance was counted for each gene, transposon, and cluster (RPKM: Reads Per Kilobase per Million mapped reads) using BEDTools (Version 2.27.1) (Quinlan and Hall 2010) and normalized to total number of genome mapping reads excluding rRNA mapping reads.

### Small RNA-seq Analysis

piSet_srnaseq was used to analyze small RNA sequencing data. Cutadapt (Version 1.15) (Martin 2011) was used to remove 3ʹ end adaptor. The raw small RNA-seq reads were first mapped to rRNA, miRNA hairpin, snoRNA, snRNA, and tRNA sequence allowing for no mismatches using Bowtie (Version 1.1.0) (B Langmead et al. 2009). The remaining reads were then mapped to the *Drosophila* genome (*dm6*) and transposon consensus sequences. Small RNA abundance was normalized to uniquely mapping reads. For Ping-Pong analysis, the 5ʹ go 5ʹ overlaps between piRNAs mapping to opposite genomic strands were calculated, and Z-score for 10 nucleotide (nt) overlap was calculated by using 1-9 nt and 11-30 nt overlaps as background (Li et al. 2009).

### Cut and Run-seq Analysis

Cut and run sequencing analysis was performed using piSet_chipseq pipeline on GitHub. Briefly, Bowtie2 (Version 2.2.5) with default settings was used to map sequencing reads to the *Drosophila* genome (*dm6*) and transposon consensus sequence (Ben Langmead and Salzberg 2012). Signal mapping to transposons were normalized to total number of genome mapping Cut and Run reads.

## Notes

### Competing Interest Statement

The authors have declared no competing interest.

## References

Ahmad, Kami. 2018. “CUT&RUN with Drosophila Tissues.” Protocols.Io, February.

Akkouche, Abdou, Bruno Mugat, Bridlin Barckmann, Carolina Varela-Chavez, Blaise Li, Raoul Raffel, Alain Pélisson, and Séverine Chambeyron. 2017. “Piwi Is Required during Drosophila Embryogenesis to License Dual-Strand PiRNA Clusters for Transposon Repression in Adult Ovaries.” Molecular Cell 66 (3): 411–419.e4. https://doi.org/10.1016/j.molcel.2017.03.017.

Bao, Weidong, Kenji K Kojima, and Oleksiy Kohany. 2015. “Repbase Update, a Database of Repetitive Elements in Eukaryotic Genomes.” Mobile DNA 6: 11. https://doi.org/10.1186/s13100-015-0041-9.

Belancio, Victoria P, Prescott L Deininger, and Astrid M Roy-Engel. 2009. “LINE Dancing in the Human Genome: Transposable Elements and Disease.” Genome Medicine 1 (10): 97. https://doi.org/10.1186/gm97.

Belgnaoui, S Mehdi, Roger G Gosden, O John Semmes, and Abdelali Haoudi. 2006. “Human LINE-1 Retrotransposon Induces DNA Damage and Apoptosis in Cancer Cells.” Cancer Cell International 6: 13. https://doi.org/10.1186/1475-2867-6-13.

Brennecke, Julius, Alexei A. Aravin, Alexander Stark, Monica Dus, Manolis Kellis, Ravi Sachidanandam, and Gregory J. Hannon. 2007. “Discrete Small RNA-Generating Loci as Master Regulators of Transposon Activity in Drosophila.” Cell 128 (6): 1089–1103. https://doi.org/10.1016/j.cell.2007.01.043.

Brennecke, Julius, Colin D. Malone, Alexei A. Aravin, Ravi Sachidanandam, Alexander Stark, and Gregory J. Hannon. 2008. “An Epigenetic Role for Maternally Inherited PiRNAs in Transposon Silencing.” *Science (New York*, N.Y.) 322 (5906): 1387–92. https://doi.org/10.1126/science.1165171.

Brewer-Jensen, Paul, Carrie B. Wilson, John Abernethy, Lonna Mollison, Samantha Card, and Lillie L. Searles. 2016. “Suppressor of Sable [Su(s)] and Wdr82 down-Regulate RNA from Heat-Shock-Inducible Repetitive Elements by a Mechanism That Involves Transcription Termination.” RNA 22 (1): 139–54. https://doi.org/10.1261/rna.048819.114.

Cappucci, Ugo, Fabrizia Noro, Assunta Maria Casale, Laura Fanti, Maria Berloco, Angela Alessandra Alagia, Luigi Grassi, Loredana Le Pera, Lucia Piacentini, and Sergio Pimpinelli. 2019. “The Hsp70 Chaperone Is a Major Player in Stress-Induced Transposable Element Activation.” Proceedings of the National Academy of Sciences of the United States of America 116 (36): 17943–50. https://doi.org/10.1073/pnas.1903936116.

Chang, Timothy H, Eugenio Mattei, Ildar Gainetdinov, Cansu Colpan, Zhiping Weng, and Phillip D Zamore. 2019. “Maelstrom Represses Canonical Polymerase II Transcription within Bi-Directional PiRNA Clusters in Drosophila Melanogaster.” Molecular Cell 73 (2): 291–303.e6. https://doi.org/10.1016/j.molcel.2018.10.038.

Chen, Yung A Chia Ariel, Evelyn Stuwe, Yicheng Luo, Maria Ninova, Adrien le Thomas, Ekaterina Rozhavskaya, Sisi Li, et al. 2016. “Cutoff Suppresses RNA Polymerase II Termination to Ensure Expression of PiRNA Precursors.” Molecular Cell 63 (1): 97–109. https://doi.org/10.1016/j.molcel.2016.05.010.

Cook, H A, B S Koppetsch, J Wu, and W E Theurkauf. 2004. “The Drosophila SDE3 Homolog Armitage Is Required for Oskar MRNA Silencing and Embryonic Axis Specification.” Cell 116 (6): 817–29. https://doi.org/10.1016/s0092-8674(04)00250-8.

Dobin, Alexander, Carrie A Davis, Felix Schlesinger, Jorg Drenkow, Chris Zaleski, Sonali Jha, Philippe Batut, Mark Chaisson, and Thomas R Gingeras. 2013. “STAR: Ultrafast Universal RNA-Seq Aligner.” *Bioinformatics (Oxford*, England) 29 (1): 15–21. https://doi.org/10.1093/bioinformatics/bts635.

ElMaghraby, M F, P R Andersen, F Puhringer, U Hohmann, K Meixner, T Lendl, L Tirian, and J Brennecke. 2019. “A Heterochromatin-Specific RNA Export Pathway Facilitates PiRNA Production.” Cell 178 (4): 964–979 e20. https://doi.org/10.1016/j.cell.2019.07.007.

Fanti, Laura, Lucia Piacentini, Ugo Cappucci, Assunta M. Casale, and Sergio Pimpinelli. 2017. “Canalization by Selection of de Novo Induced Mutations.” Genetics 206 (4): 1995–2006. https://doi.org/10.1534/genetics.117.201079.

Fast, Isabel, and David Rosenkranz. 2018. “Temperature-Dependent Small RNA Expression in Drosophila Melanogaster.” RNA Biology 15 (3): 308–13. https://doi.org/10.1080/15476286.2018.1429881.

Feschotte, Cédric, and Ellen J Pritham. 2007. “DNA Transposons and the Evolution of Eukaryotic Genomes.” Annual Review of Genetics 41 (6): 331–68. https://doi.org/10.1146/annurev.genet.40.110405.090448.

Findley, Seth D, Mio Tamanaha, Nigel J Clegg, and Hannele Ruohola-Baker. 2003. “Maelstrom, a Drosophila Spindle-Class Gene, Encodes a Protein That Colocalizes with Vasa and RDE1/AGO1 Homolog, Aubergine, in Nuage.” *Development (Cambridge*, England) 130 (5): 859–71. https://doi.org/10.1242/dev.00310.

Funikov, S. Yu, SS S Ryazansky, ES S Zelentsova, VI I Popenko, OG G Leonova, DG G Garbuz, MB B Evgen’ev, and OG G Zatsepina. 2015. “The Peculiarities of PiRNA Expression upon Heat Shock Exposure in Drosophila Melanogaster.” Mobile Genetic Elements 5 (5): 72–80. https://doi.org/10.1080/2159256X.2015.1086502.

Garrigues, Jacob M, Brian V Tsu, Matthew D Daugherty, and Amy E Pasquinelli. 2019. “Diversification of the Caenorhabditis Heat Shock Response by Helitron Transposable Elements.” ELife 8 (December). https://doi.org/10.7554/eLife.51139.

Ge, Daniel Tianfang, Wei Wang, Cindy Tipping, Ildar Gainetdinov, Zhiping Weng, and Phillip D Zamore. 2019. “The RNA-Binding ATPase, Armitage, Couples PiRNA Amplification in Nuage to Phased PiRNA Production on Mitochondria.” Molecular Cell 74 (5): 982–995.e6. https://doi.org/10.1016/j.molcel.2019.04.006.

Gebert, Daniel, Lena K. Neubert, Catrin Lloyd, Jinghua Gui, Ruth Lehmann, and Felipe Karam Teixeira. 2021. “Large Drosophila Germline PiRNA Clusters Are Evolutionarily Labile and Dispensable for Transposon Regulation.” Molecular Cell 81 (19): 3965–3978.e5. https://doi.org/10.1016/j.molcel.2021.07.011.

Grandbastien, M A, H Lucas, J B Morel, C Mhiri, S Vernhettes, and J M Casacuberta. 1997. “The Expression of the Tobacco Tnt1 Retrotransposon Is Linked to Plant Defense Responses.” Genetica 100 (1–3): 241–52. http://www.ncbi.nlm.nih.gov/pubmed/9440277.

Han, Bo W., Wei Wang, Chengjian Li, Zhiping Weng, and Phillip D. Zamore. 2015. “PiRNA-Guided Transposon Cleavage Initiates Zucchini-Dependent, Phased PiRNA Production.” Science 348 (6236): 817–21. https://doi.org/10.1126/science.aaa1264.

Harris, a N, and P M Macdonald. 2001. “Aubergine Encodes a Drosophila Polar Granule Component Required for Pole Cell Formation and Related to EIF2C.” *Development (Cambridge*, England) 128 (14): 2823–32.

Hur, Junho K, and Yun Doo Chung. 2016. “A Novel Model of THO/TREX Loading onto Target RNAs in Metazoan Gene Expression.” BMB Reports 49 (7): 355–56. https://doi.org/10.1101/gad.276030.115.

Hur, Junho K., Yicheng Luo, Sungjin Moon, Maria Ninova, Georgi K. Marinov, Yun D. Chung, and Alexei A. Aravin. 2016. “Splicing-Independent Loading of TREX on Nascent RNA Is Required for Efficient Expression of Dual-Strand PiRNA Clusters in Drosophila.” Genes and Development 30 (7): 840–55. https://doi.org/10.1101/gad.276030.115.

Jamrich, M, A L Greenleaf, and E K Bautz. 1977. “Localization of RNA Polymerase in Polytene Chromosomes of Drosophila Melanogaster.” Proceedings of the National Academy of Sciences of the United States of America 74 (5): 2079–83. https://doi.org/10.1073/pnas.74.5.2079.

Jardim, Sinara Santos, André Passaglia Schuch, Camila Moura Pereira, and Elgion Lucio Silva Loreto. 2015. “Effects of Heat and UV Radiation on the Mobilization of Transposon Mariner-Mos1.” Cell Stress & Chaperones 20 (5): 843–51. https://doi.org/10.1007/s12192-015-0611-2.

Kelleher, Erin S, Jyoti Lama, and Luyang Wang. 2020. “Uninvited Guests: How Transposable Elements Take Advantage of Drosophila Germline Stem Cells, and How Stem Cells Fight Back.” Current Opinion in Insect Science 37 (February): 49–56. https://doi.org/10.1016/j.cois.2019.11.002.

Kim, Daehwan, Ben Langmead, and Steven L Salzberg. 2015. “HISAT: A Fast Spliced Aligner with Low Memory Requirements.” Nature Methods 12 (4): 357–60. https://doi.org/10.1038/nmeth.3317.

Klattenhoff, Carla, Hualin Xi, Chengjian Li, Soohyun Lee, Jia Xu, Jaspreet S. Khurana, Fan Zhang, et al. 2009. “The Drosophila HP1 Homolog Rhino Is Required for Transposon Silencing and PiRNA Production by Dual-Strand Clusters.” Cell 138 (6): 1137–49. https://doi.org/10.1016/j.cell.2009.07.014.

Klenov, Mikhail S, Olesya A Sokolova, Evgeny Y Yakushev, Anastasia D Stolyarenko, Elena A Mikhaleva, Sergey A Lavrov, and Vladimir A Gvozdev. 2011. “Separation of Stem Cell Maintenance and Transposon Silencing Functions of Piwi Protein.” Proceedings of the National Academy of Sciences of the United States of America 108 (46): 18760–65. https://doi.org/10.1073/pnas.1106676108.

Kneuss, Emma, Marzia Munafò, Evelyn L Eastwood, Undine-Sophie Deumer, Jonathan B Preall, Gregory J Hannon, and Benjamin Czech. 2019. “Specialization of the Drosophila Nuclear Export Family Protein Nxf3 for PiRNA Precursor Export.” Genes & Development 33 (17–18): 1208–20. https://doi.org/10.1101/gad.328690.119.

Kuan, Yung-Shu, Paul Brewer-Jensen, Wen-Li Bai, Cedric Hunter, Carrie B. Wilson, Sarah Bass, John Abernethy, James S. Wing, and Lillie L. Searles. 2009. “Drosophila Suppressor of Sable Protein [Su(s)] Promotes Degradation of Aberrant and Transposon-Derived RNAs.” Molecular and Cellular Biology 29 (20): 5590–5603. https://doi.org/10.1128/mcb.00039-09.

Langmead, B, C Trapnell, M Pop, and S L Salzberg. 2009. “Ultrafast and Memory-Efficient Alignment of Short DNA Sequences to the Human Genome.” Genome Biol 10 (3): R25. https://doi.org/10.1186/gb-2009-10-3-r25.

Langmead, Ben, and Steven L Salzberg. 2012. “Fast Gapped-Read Alignment with Bowtie 2.” Nature Methods 9 (4): 357–59. https://doi.org/10.1038/nmeth.1923.

Li, Chengjian, Vasily V Vagin, Soohyun Lee, Jia Xu, Shengmei Ma, Hualin Xi, Hervé Seitz, et al. 2009. “Collapse of Germline PiRNAs in the Absence of Argonaute3 Reveals Somatic PiRNAs in Flies.” Cell 137 (3): 509–21. https://doi.org/10.1016/j.cell.2009.04.027.

Lim, Ai Khim, and Toshie Kai. 2007. “Unique Germ-Line Organelle, Nuage, Functions to Repress Selfish Genetic Elements in Drosophila Melanogaster.” Proceedings of the National Academy of Sciences of the United States of America 104 (16): 6714–19. https://doi.org/10.1073/pnas.0701920104.

Lin, H, and A C Spradling. 1997. “A Novel Group of Pumilio Mutations Affects the Asymmetric Division of Germline Stem Cells in the Drosophila Ovary.” *Development (Cambridge*, England) 124 (12): 2463–76. https://doi.org/10.1242/dev.124.12.2463.

Lindquist, S. 1986. “The Heat-Shock Response.” Annual Review of Biochemistry 55: 1151–91. https://doi.org/10.1146/annurev.bi.55.070186.005443.

Malone, Colin D., Julius Brennecke, Monica Dus, Alexander Stark, W. Richard McCombie, Ravi Sachidanandam, and Gregory J. Hannon. 2009. “Specialized PiRNA Pathways Act in Germline and Somatic Tissues of the Drosophila Ovary.” Cell 137 (3): 522–35. https://doi.org/10.1016/j.cell.2009.03.040.

Martin, Marcel. 2011. “Cutadapt Removes Adapter Sequences from High-Throughput Sequencing Reads.” EMBnet.Journal 17 (1): 10. https://doi.org/10.14806/ej.17.1.200.

Maside, Xulio, Carolina Bartolomé, and Brian Charlesworth. 2002. “S-Element Insertions Are Associated with the Evolution of the Hsp70 Genes in Drosophila Melanogaster.” Current Biology : CB 12 (19): 1686–91. https://doi.org/10.1016/s0960-9822(02)01181-8.

McClintock, B. 1984. “The Significance of Responses of the Genome to Challenge.” *Science (New York*, N.Y.) 226 (4676): 792–801. https://doi.org/10.1126/science.15739260.

Mendel, M, and R S Pillai. 2019. “Nxf3: A Middleman with the Right Connections for Unspliced PiRNA Precursor Export.” Genes Dev 33 (17–18): 1095–97. https://doi.org/10.1101/gad.330530.119.

Mohn, Fabio, Dominik Handler, and Julius Brennecke. 2015. “Noncoding RNA. PiRNA-Guided Slicing Specifies Transcripts for Zucchini-Dependent, Phased PiRNA Biogenesis.” *Science (New York*, N.Y.) 348 (6236): 812–17. https://doi.org/10.1126/science.aaa1039.

Mohn, Fabio, Grzegorz Sienski, Dominik Handler, and Julius Brennecke. 2014. “The Rhino-Deadlock-Cutoff Complex Licenses Noncanonical Transcription of Dual-Strand PiRNA Clusters in Drosophila.” Cell 157 (6): 1364–79. https://doi.org/10.1016/j.cell.2014.04.031.

Mombach, Daniela Moreira, Tiago Minuzzi Freire da Fontoura Gomes, and Elgion Lucio Silva Loreto. 2022. “Stress Does Not Induce a General Transcription of Transposable Elements in Drosophila.” Molecular Biology Reports 49 (9): 9033–40. https://doi.org/10.1007/s11033-022-07839-7.

Moon, Sungjin, Bongki Cho, Su-Hong Min, Daekee Lee, and Yun Doo Chung. 2011. “The THO Complex Is Required for Nucleolar Integrity in Drosophila Spermatocytes.” *Development (Cambridge*, England) 138 (17): 3835–45. https://doi.org/10.1242/dev.056945.

Pane, Attilio, Kristina Wehr, and Trudi Schüpbach. 2007. “Zucchini and Squash Encode Two Putative Nucleases Required for RasiRNA Production in the Drosophila Germline.” Developmental Cell 12 (6): 851–62. https://doi.org/10.1016/j.devcel.2007.03.022.

Parhad, Swapnil S., Shikui Tu, Zhiping Weng, and William E. Theurkauf. 2017. “Adaptive Evolution Leads to Cross-Species Incompatibility in the PiRNA Transposon Silencing Machinery.” Developmental Cell 43 (1): 60–70.e5. https://doi.org/10.1016/j.devcel.2017.08.012.

Parhad, Swapnil S., Tianxiong Yu, Gen Zhang, Nicholas P. Rice, Zhiping Weng, and William E. Theurkauf. 2020. “Adaptive Evolution Targets a PiRNA Precursor Transcription Network.” Cell Reports 30 (8): 2672–2685.e5. https://doi.org/10.1016/j.celrep.2020.01.109.

Payer, Lindsay M., and Kathleen H. Burns. 2019. “Transposable Elements in Human Genetic Disease.” Nature Reviews Genetics 20 (12): 760–72. https://doi.org/10.1038/s41576-019-0165-8.

Prasad, Mohit, Anna C-C Jang, Michelle Starz-Gaiano, Mariana Melani, and Denise J Montell. 2007. “A Protocol for Culturing Drosophila Melanogaster Stage 9 Egg Chambers for Live Imaging.” Nature Protocols 2 (10): 2467–73. https://doi.org/10.1038/nprot.2007.363.

Quinlan, Aaron R, and Ira M Hall. 2010. “BEDTools: A Flexible Suite of Utilities for Comparing Genomic Features.” *Bioinformatics (Oxford*, England) 26 (6): 841–42. https://doi.org/10.1093/bioinformatics/btq033.

Raje, Himanshu S, Molly E Lieux, and Patrick J DiMario. 2018. “R1 Retrotransposons in the Nucleolar Organizers of Drosophila Melanogaster Are Transcribed by RNA Polymerase I upon Heat Shock.” Transcription 9 (5): 273–85. https://doi.org/10.1080/21541264.2018.1506682.

Rehwinkel, J, A Herold, K Gari, T Kocher, M Rode, F L Ciccarelli, M Wilm, and E Izaurralde. 2004. “Genome-Wide Analysis of MRNAs Regulated by the THO Complex in Drosophila Melanogaster.” Nat Struct Mol Biol 11 (6): 558–66. https://doi.org/10.1038/nsmb759.

Roquis, David, Marta Robertson, Liang Yu, Michael Thieme, Magdalena Julkowska, and Etienne Bucher. 2021. “Genomic Impact of Stress-Induced Transposable Element Mobility in Arabidopsis.” Nucleic Acids Research 49 (18): 10431–47. https://doi.org/10.1093/nar/gkab828.

Shpiz, Sergey, Sergei Ryazansky, Ivan Olovnikov, Yuri Abramov, and Alla Kalmykova. 2014. “Euchromatic Transposon Insertions Trigger Production of Novel Pi- and Endo-SiRNAs at the Target Sites in the Drosophila Germline.” PLoS Genetics 10 (2). https://doi.org/10.1371/journal.pgen.1004138.

Sienski, Grzegorz, Julia Batki, Kirsten-andré Senti, Derya Dönertas, Laszlo Tirian, Katharina Meixner, and Julius Brennecke. 2015. “Silencio / CG9754 Connects the Piwi – PiRNA Complex to the Cellular Heterochromatin Machinery.” Genes and Development 29: 1–14. https://doi.org/10.1101/gad.271908.115.

Sienski, Grzegorz, Derya Dönertas, and Julius Brennecke. 2012. “Transcriptional Silencing of Transposons by Piwi and Maelstrom and Its Impact on Chromatin State and Gene Expression.” Cell 151 (5): 964–80. https://doi.org/10.1016/j.cell.2012.10.040.

Skene, Peter J, and Steven Henikoff. 2017. “An Efficient Targeted Nuclease Strategy for High-Resolution Mapping of DNA Binding Sites.” ELife 6 (January). https://doi.org/10.7554/eLife.21856.

Styhler, S, Akira Nakamura, A Swan, B Suter, and Paul Lasko. 1998. “Vasa Is Required for GURKEN Accumulation in the Oocyte, and Is Involved in Oocyte Differentiation and Germline Cyst Development.” Development 125 (9): 1569–78. https://doi.org/10.1242/dev.125.9.1569.

Sun, Linhua, Yuqing Jing, Xinyu Liu, Qi Li, Zhihui Xue, Zhukuan Cheng, Daowen Wang, Hang He, and Weiqiang Qian. 2020. “Heat Stress-Induced Transposon Activation Correlates with 3D Chromatin Organization Rearrangement in Arabidopsis.” Nature Communications 11 (1): 1886. https://doi.org/10.1038/s41467-020-15809-5.

Takehira, Kanako, Yui Hayashi, Kosuke Nozawa, Lu Chen, Takamasa Suzuki, Yukari Masuta, Atsushi Kato, and Hidetaka Ito. 2021. “DRD1, a SWI/SNF-like Chromatin Remodeling Protein, Regulates a Heat-Activated Transposon in Arabidopsis Thaliana.” Genes & Genetic Systems 96 (3): 151–58. https://doi.org/10.1266/ggs.21-00005.

Teves, Sheila S, and Steven Henikoff. 2011. “Heat Shock Reduces Stalled RNA Polymerase II and Nucleosome Turnover Genome-Wide.” Genes & Development 25 (22): 2387–97. https://doi.org/10.1101/gad.177675.111.

Theurkauf, W E. 1994. “Immunofluorescence Analysis of the Cytoskeleton during Oogenesis and Early Embryogenesis.” Methods Cell Biol 44: 489–505. https://doi.org/10.1016/s0091-679x(08)60928-0.

Vagin, Vasily v., Yang Yu, Anna Jankowska, Yicheng Luo, Kaja A. Wasik, Colin D. Malone, Emily Harrison, et al. 2013. “Minotaur Is Critical for Primary PiRNA Biogenesis.” *RNA (New York*, N.Y.) 19 (8): 1064–77. https://doi.org/10.1261/rna.039669.113.

Volpe, Alison M, Heidi Horowitz, Constance M Grafer, Stephen M Jackson, and Celeste A Berg. 2001. “Drosophila *Rhino* Encodes a Female-Specific Chromo-Domain Protein That Affects Chromosome Structure and Egg Polarity.” Genetics 159 (3): 1117–34. https://doi.org/10.1093/genetics/159.3.1117.

Wang, W, B W Han, C Tipping, D T Ge, Z Zhang, Z Weng, and P D Zamore. 2015. “Slicing and Binding by Ago3 or Aub Trigger Piwi-Bound PiRNA Production by Distinct Mechanisms.” Mol Cell 59 (5): 819–30. https://doi.org/10.1016/j.molcel.2015.08.007.

Webster, Alexandre, Sisi Li, Junho K Hur, Malte Wachsmuth, Justin S Bois, Edward M Perkins, Dinshaw J Patel, and Alexei A Aravin. 2015. “Aub and Ago3 Are Recruited to Nuage through Two Mechanisms to Form a Ping-Pong Complex Assembled by Krimper.” Molecular Cell 59 (4): 564–75. https://doi.org/10.1016/j.molcel.2015.07.017.

Xiol, Jordi, Pietro Spinelli, Maike A. Laussmann, David Homolka, Zhaolin Yang, Elisa Cora, Yohann Coute, et al. 2014. “RNA Clamping by Vasa Assembles a PiRNA Amplifier Complex on Transposon Transcripts.” Cell 157 (7): 1698– 1711. https://doi.org/10.1016/j.cell.2014.05.018.

Yu, Tianxiong, Birgit S Koppetsch, Sara Pagliarani, Stephen Johnston, Noah J Silverstein, Jeremy Luban, Keith Chappell, Zhiping Weng, and William E Theurkauf. 2019. “The PiRNA Response to Retroviral Invasion of the Koala Genome.” Cell 179 (3): 632–643.e12. https://doi.org/10.1016/j.cell.2019.09.002.

Yu, Yang, Jiaqi Gu, Ying Jin, Yicheng Luo, Jonathan B. Preall, Jinbiao Ma, Benjamin Czech, and Gregory J. Hannon. 2015. “Panoramix Enforces PiRNA-Dependent Cotranscriptional Silencing.” Science 350 (6258): 339–42. https://doi.org/10.1126/science.aab0700.

Zhang, Fan, Jie Wang, Jia Xu, Zhao Zhang, Birgit S Koppetsch, Nadine Schultz, Thom Vreven, et al. 2012. “UAP56 Couples PiRNA Clusters to the Perinuclear Transposon Silencing Machinery.” Cell 151 (4): 871–84. https://doi.org/10.1016/j.cell.2012.09.040.

Zhang, G, S Tu, T Yu, X O Zhang, S S Parhad, Z Weng, and W E Theurkauf. 2018. “Co-Dependent Assembly of Drosophila PiRNA Precursor Complexes and PiRNA Cluster Heterochromatin.” Cell Rep 24 (13): 3413–3422 e4. https://doi.org/10.1016/j.celrep.2018.08.081.

Zhang, Gen, Tianxiong Yu, Swapnil S Parhad, Samantha Ho, Zhiping Weng, and William E Theurkauf. 2021. “PiRNA-Independent Transposon Silencing by the Drosophila THO Complex.” Developmental Cell 56 (18): 2623–2635.e5. https://doi.org/10.1016/j.devcel.2021.08.021.

Zhang, Zhao, Jie Wang, Nadine Schultz, Fan Zhang, Swapnil S Parhad, Shikui Tu, Thom Vreven, Phillip D Zamore, Zhiping Weng, and William E Theurkauf. 2014. “The HP1 Homolog Rhino Anchors a Nuclear Complex That Suppresses PiRNA Precursor Splicing.” Cell 157 (6): 1353–63. https://doi.org/10.1016/j.cell.2014.04.030.

