## Supplemental Figures for "Rapid disassembly and Piwi-independent reassembly of *Drosophila* piRNA cluster heterochromatin following acute heat shock"

#### Supplemental Figure Legends

**Figure S1: Cluster expression, but not genes or piRNAs, decrease with heat shock**

A. Density scatter plots comparing gene expression (rpkm, log10 scale) in *w¹* control vs. samples heat shocked at 37°C for 1 hour and allowed to recover for 1, 12, and 24 hours. Red dots are the HSP genes. Dotted line is a 2-fold change.

B. All mapping long RNA signal tracks for cluster 42AB in *w¹* control, w1 heat shocked for 1 hour at 37°C allowed to recover for 1 hour, 12 hours, and 24 hours. Heat shocked samples show overexpression at embedded elements maximally at 1 hour, and this expression decreases with time.

C. Scatter plot comparing all anti-sense piRNAs mapping to transposon families (rpkm, log10 scale), piRNAs uniquely mapping to clusters (rpkm, log10 scale), and piRNAs Zscore (log2 scale), a measure of the enrichment of ping-pong piRNA pairs, in *w¹* control vs. samples heat shocked at 37°C for 1 hour and allowed to recover for 1 hour. The dotted line is a 2-fold change. The red dot is the major cluster 42AB.

D. Scatter plot comparing unique mapping piRNA clusters expression (rpkm, log10 scale) in *w¹* control vs. samples heat shocked at 37°C for 1 hour and allowed to recover for 1, 12, and 24 hours. The dotted line is 2-fold change, the red dot is the major cluster 42AB. Most clusters show a trend towards reduced expression with some recovery with time.

**Figure S2: Absolute temperature matters in displacing Rhino foci**

A. Live confocal images of GFP-Rhi dissected egg chambers heat shocked for 1 hour for the indicated temperature compared to control. Heat shock at 34°C shows no loss of

Rhino foci, whereas 35°C loses Rhino localization 1 Hour after heat shock at 35°C Rhino foci return. The area highlighted in the box is a digitally zoomed-in single nurse cell nuclei.

B. Live confocal images of GFP-Rhi dissected egg chambers from flies raised at 18°C heat shocked for 1 hour at 30°C and control. Heat shock at 30°C shows no loss of Rhino foci. The area highlighted in the box is a digitally zoomed-in single nurse cell nuclei.

**Figure S3: Disassembly and reassembly of other piRNA precursor proteins involved in transcription and export.**

A. Confocal images of a nurse cell nucleus in GFP-Cuff control and heat shocked at 37°C for 30 minutes labeled for Deadlock (anti-Del). Arrows point at Del foci in control.

B. Scatterplot comparing the normalized signal of Rhino and GPF-Cuff foci heat shock at 37°C for an hour and allowed to recover for 3 hours and 45 mins. A single nurse cell nucleus with visible Rhino staining was selected to be the only nucleus in the optical section and then imaged in 1-micrometer steps. A minimum of four nuclei was used for each sample. Each point represents a focus of Rhino or GFP-Cuff five standard deviations above the background with the signal normalized to the maximum fluorescence in control. Rhino and GFP-Cuff colocalize similarly to no shock control indicating that Rhino and Cuff recover quickly. Note that most nuclei as this time point have no Rhino foci (See Figure 3J)

C. Confocal images of a nurse cell nucleus in GFP-Cuff from above labeled for GFP-Cuff (anti-GFP, Red) Rhino (anti-Rhino, Green) and merge with arrows pointing at the inset, which shows colocalized GFP-Cuff and Rhino signal.

D. Confocal images of an egg chamber in Venus-UAP56 labeled for Venus-UAP56 (anti-GFP, Red) Rhino (anti-Rhino, Green) and merge with arrows pointing at the inset. Control shows colocalized Venus-UAP56 and Rhino signal. One hour after a heat shock at 37$^{o}$C for 1 hour, only Venus-UAP56 signal is visible. Six hours after heat shock, colocalized Venus-UAP56 and Rhino signal is visible top inset with a clear arrow, and Rhino signal without UAP56 is also visible (white arrow, lower inset).

**Figure S4: Heat Shock alters the localization of two other piRNA pathway proteins, Squash and Maelstrom.**

A. Confocal images of a $w^1$ egg chamber stained for the piRNA pathway proteins Squash (anti-squash, green) and Ago3 (anti-ago3, red) in control and heat shocked at 37$^{o}$C for 1 hour with 1 hour recovery. Squash loses localization to nuage, while Ago3 appears unaffected.

B. Confocal images of a GFP-Maelstrom egg chamber stained for anti-GFP in control and heat shocked at 37$^{o}$C for 1 hour with 1 hour recovery. Maelstrom is generally cytoplasmic in control, and this cytoplasmic signal is confined to the nuclear periphery after heat shock

C. Confocal images of a $w^1$ egg chamber stained for the piRNA pathway protein Armi GFP in control and heat shocked at 37$^{o}$C for 1 hour with 1 hour recovery. Armi shows both nuage and mitochondrial staining in both control and after heat shock.

**Figure S5: Mutants in *panx*,, and *uap56* do not affect the speed of Rhino foci recovery**

A. Confocal images of egg chambers stained for Rhino in $panx^{M/dDf}$ mutants in control, and 2, 4, and 6 hours after a 1 hour 37$^{o}$C heat shock. For clarity, circles show outlines of nurse cell nuclei in the field of view. Two hours after heat shock, nuclei show no Rhino foci. Some nuclei show a few Rhino foci four hours after heat shock, and multiple Rhino foci are visible six hours after heat shock.

B. Confocal images of egg chambers stained for Rhino in *uap56*$^{sz15/28}$ mutants in control, and 2, 4, and 6 hours after a 1 hour 37$^{o}$C heat shock. For clarity, circles show outlines of nurse cell nuclei in the field of view. Two and four hours after heat shock, nuclei show no Rhino foci. Six hours after heat shock, multiple Rhino foci are visible.

**Figure S6: Maternal 42AB piRNAs are not required for licensing rhino binding in *w*$^{1}$, and multi-mapping piRNAs do not generate Rhino binding.**

A. Experimental Crosses using *w*$^{1}$ as the wildtype strain crossed to 42AB Df in both directions to generate genetically identical F1 that differ only in the pool of maternal piRNAs inherited.

B. Unique mapping small RNA reads mapping to 42AB cluster from the above F1 with and without uniquely mapping maternal piRNAs against 42AB Cluster

C. Cut and Run signal from the above F1 for IgG control and Rhino at 42AB cluster.

D. Example scatter plot comparing unique piRNA expression (rpkm log10) for each piRNA cluster between the two F1 progeny with and without maternal 42AB piRNAs.

E. Example scatter plot comparing all mapping piRNA expression (rpkm log10) for each transposon family between the two F1 progeny with and without maternal 42AB piRNAs.

F. Genome browser view of the repetitive genomic region surrounding the alpha gamma-element. Tracks display multi mapping piRNAs, unique mapping piRNAs and Cut and Run signal for ♀OreR x ♂42ABDf F1 (+/42AB Df) and 42ABDf x ♂OreR F1

(42AB Df/+). Both F1 populations show abundant multi-mapping piRNAs with little uniquely mapping piRNAs to this region. Despite the abundance of multi-mapping piRNAs, Cut and Run experiments for Rhino show no enrichment for this region relative to IgG controls.

### Figure S1

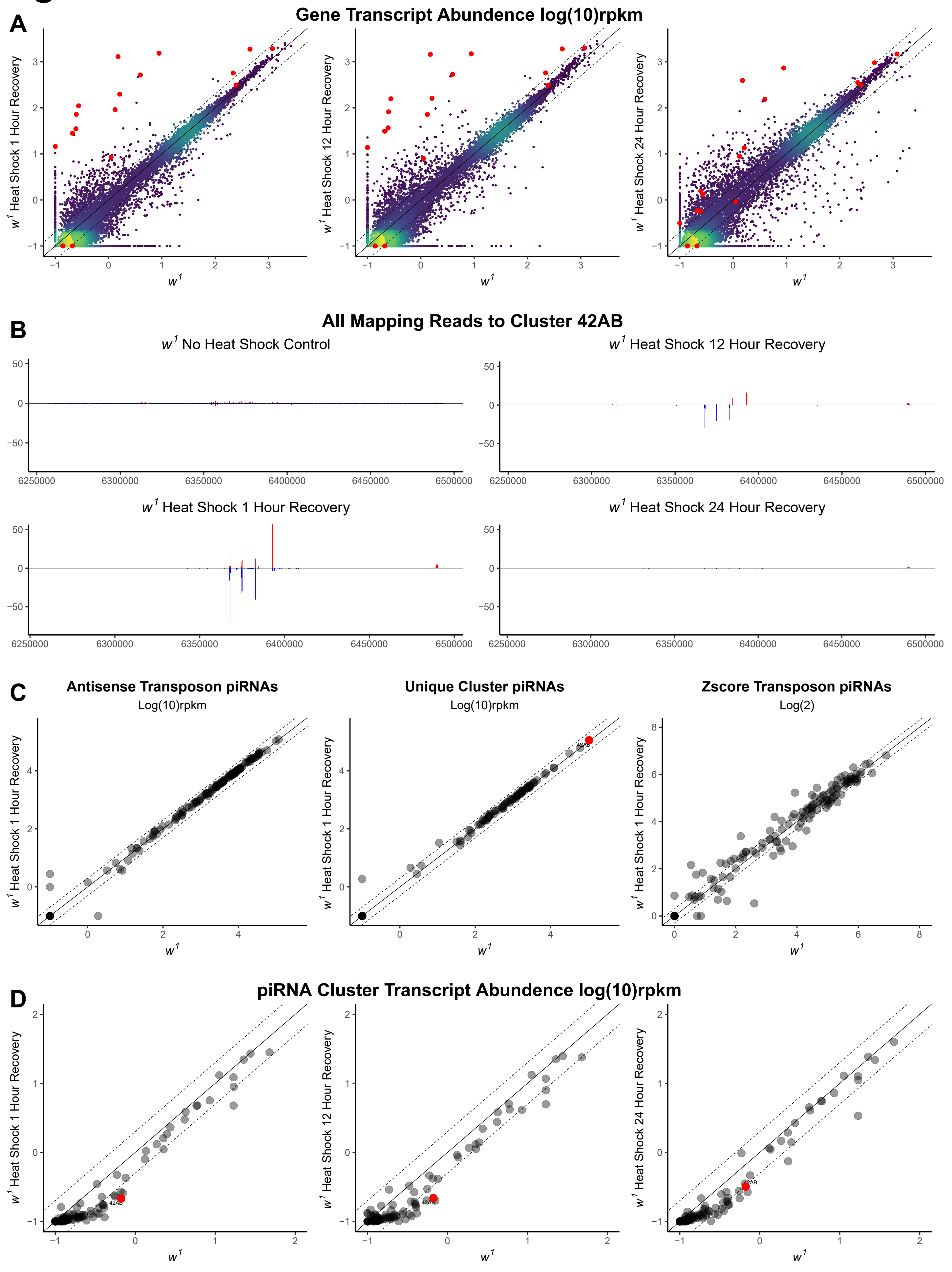
A
Gene Transcript Abundence log(10)rpkm
w1 Heat Shock 1 Hour Recovery
w1 Heat Shock 12 Hour Recovery
w1 Heat Shock 24 Hour Recovery
w1
B
All Mapping Reads to Cluster 42AB
w1 No Heat Shock Control
w1 Heat Shock 12 Hour Recovery
w1 Heat Shock 1 Hour Recovery
w1 Heat Shock 24 Hour Recovery
6250000
6300000
6350000
6400000
6450000
6500000
C
Antisense Transposon piRNAs
Log(10)rpkm
Unique Cluster piRNAs
Log(10)rpkm
Zscore Transposon piRNAs
Log(2)
42AB
D
piRNA Cluster Transcript Abundence log(10)rpkm
42AB

### Figure S2

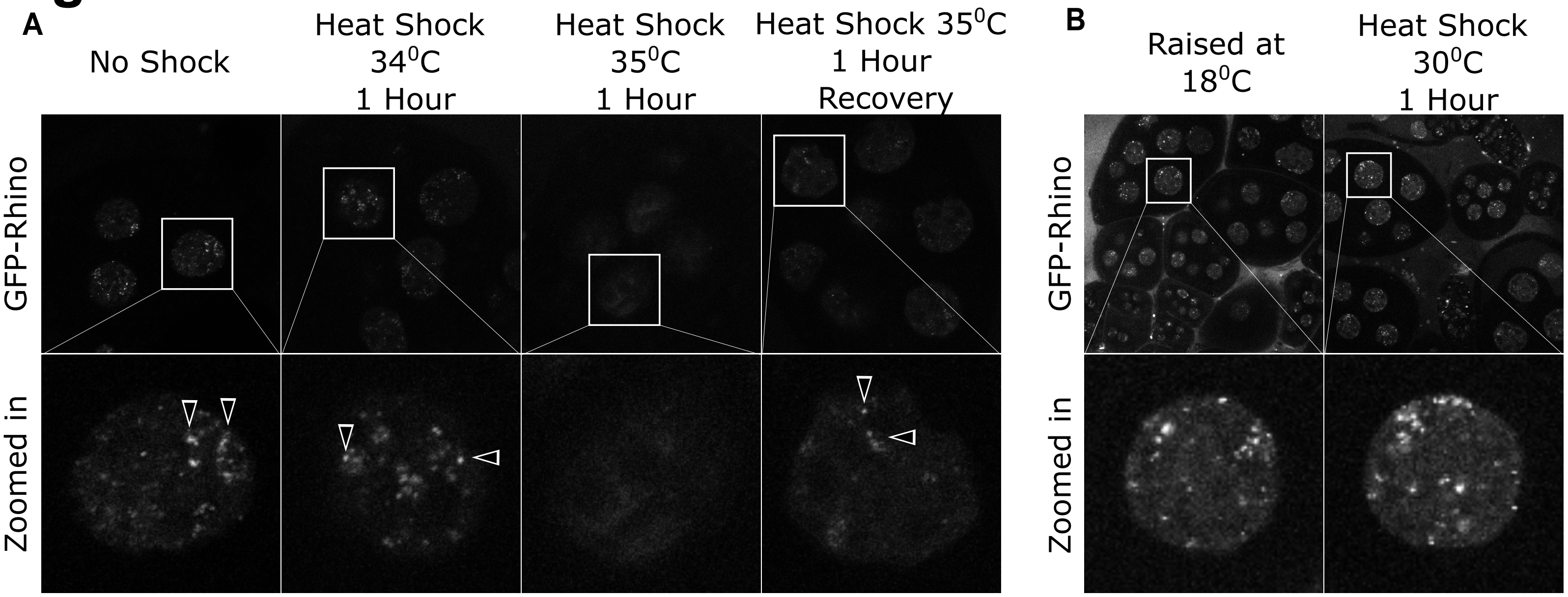

A
No Shock
Heat Shock 34⁰C 1 Hour
Heat Shock 35⁰C 1 Hour
Heat Shock 35⁰C 1 Hour Recovery
B
Raised at 18⁰C
Heat Shock 30⁰C 1 Hour
GFP-Rhino
Zoomed in

### Figure S3

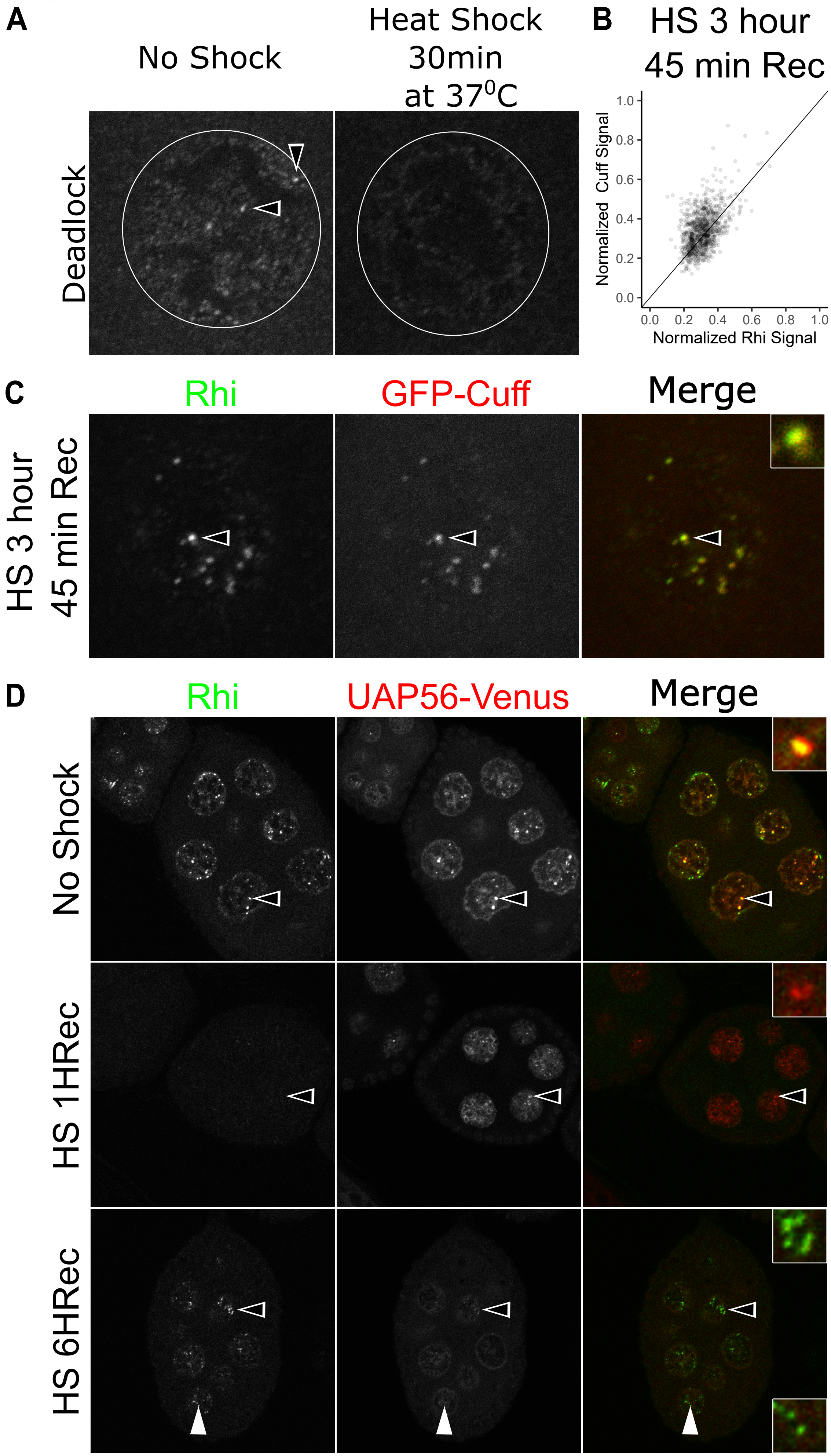
A
No Shock
Heat Shock 30min at 37$^0$C
Deadlock
B
HS 3 hour 45 min Rec
Normalized Cuff Signal
Normalized Rhi Signal
0.0
0.2
0.4
0.6
0.8
1.0
C
Rhi
GFP-Cuff
Merge
HS 3 hour 45 min Rec
D
Rhi
UAP56-Venus
Merge
No Shock
HS 1HRec
HS 6HRec

Figure S4

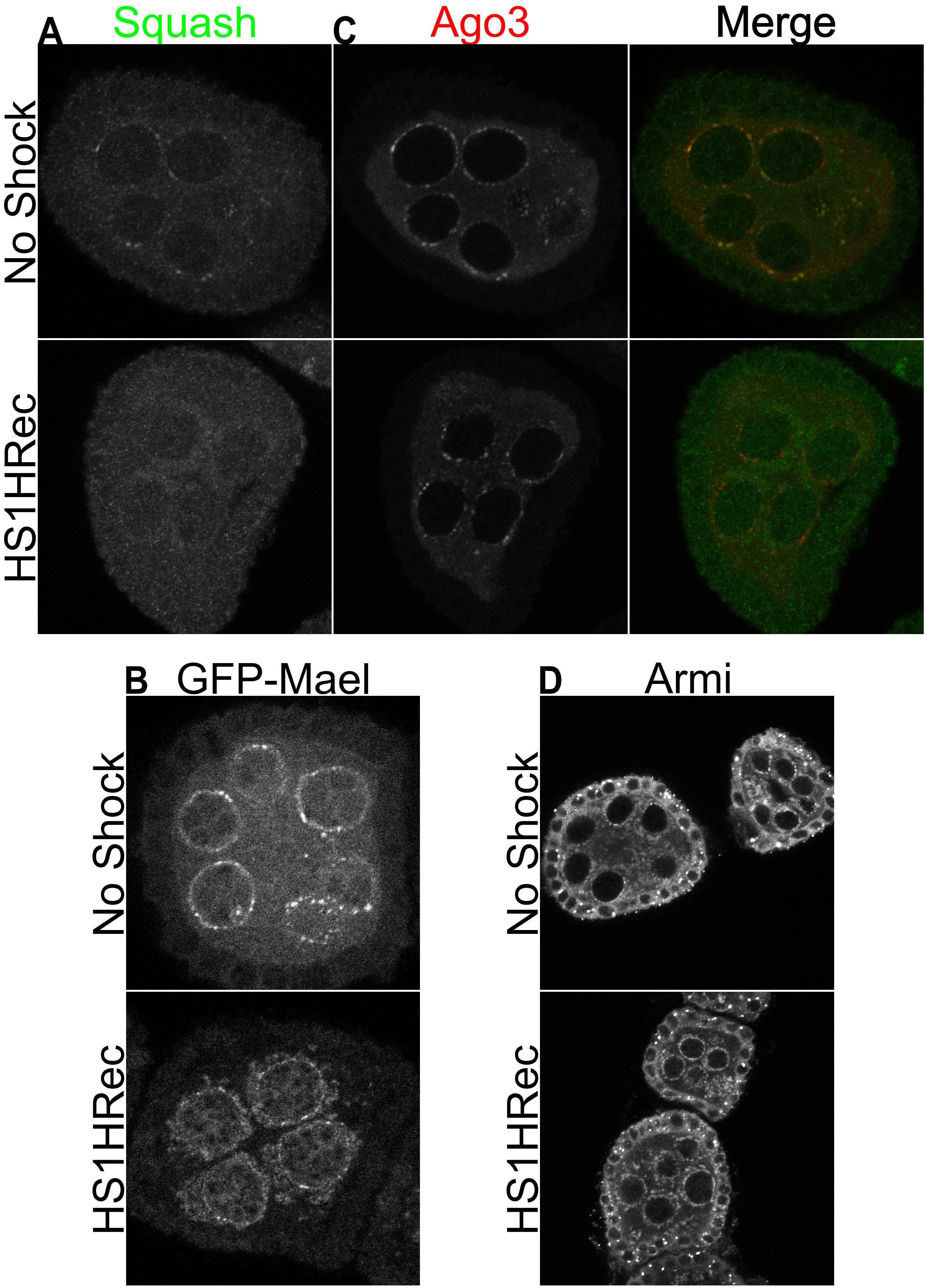
A
Squash
C
Ago3
Merge
No Shock
HS1HRec
B
GFP-Mael
D
Armi
No Shock
HS1HRec

### Figure S5

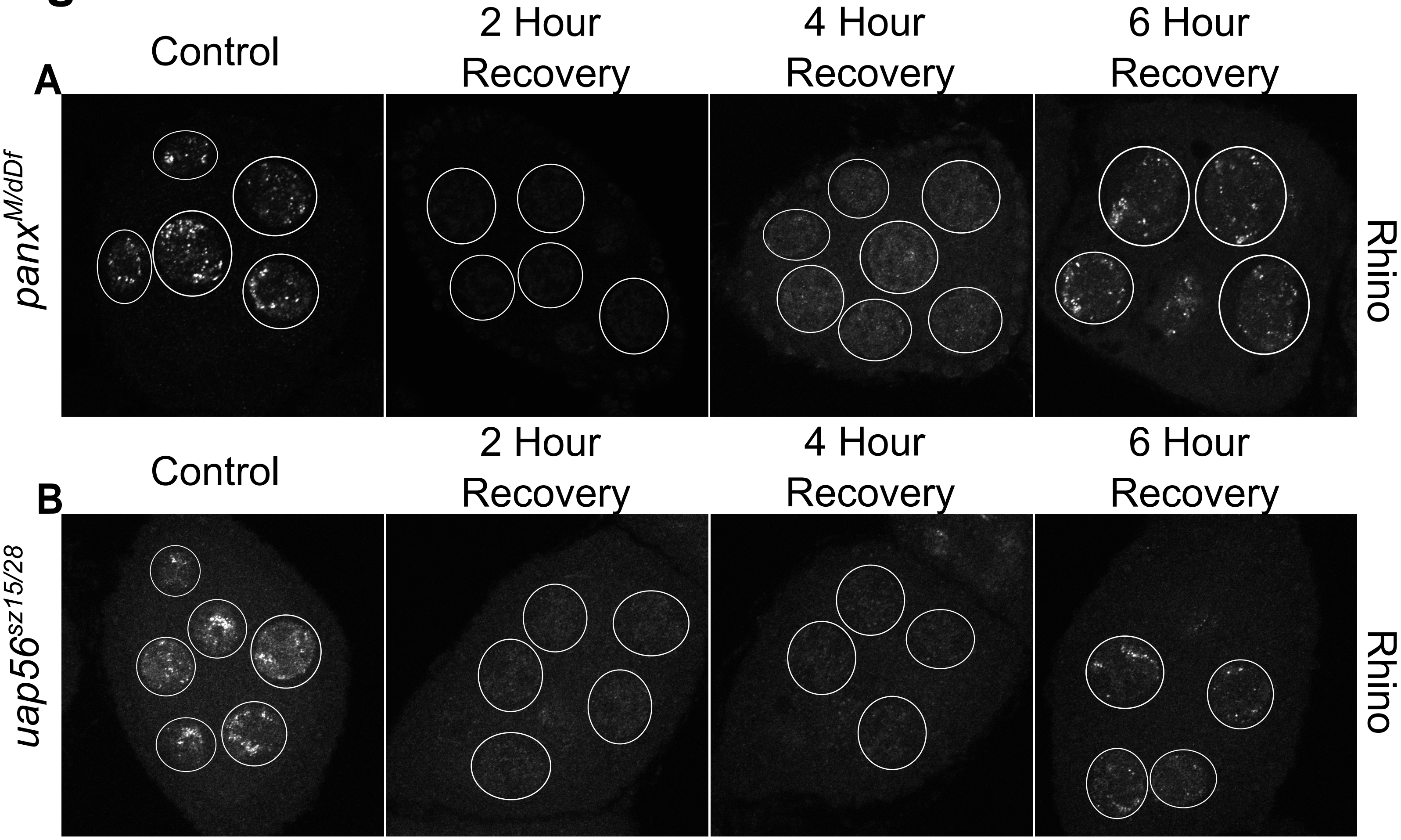
A
Control
2 Hour Recovery
4 Hour Recovery
6 Hour Recovery
panx$^{M/dDf}$
Rhino
B
Control
2 Hour Recovery
4 Hour Recovery
6 Hour Recovery
uap56$^{sz15/28}$
Rhino

### Figure S6

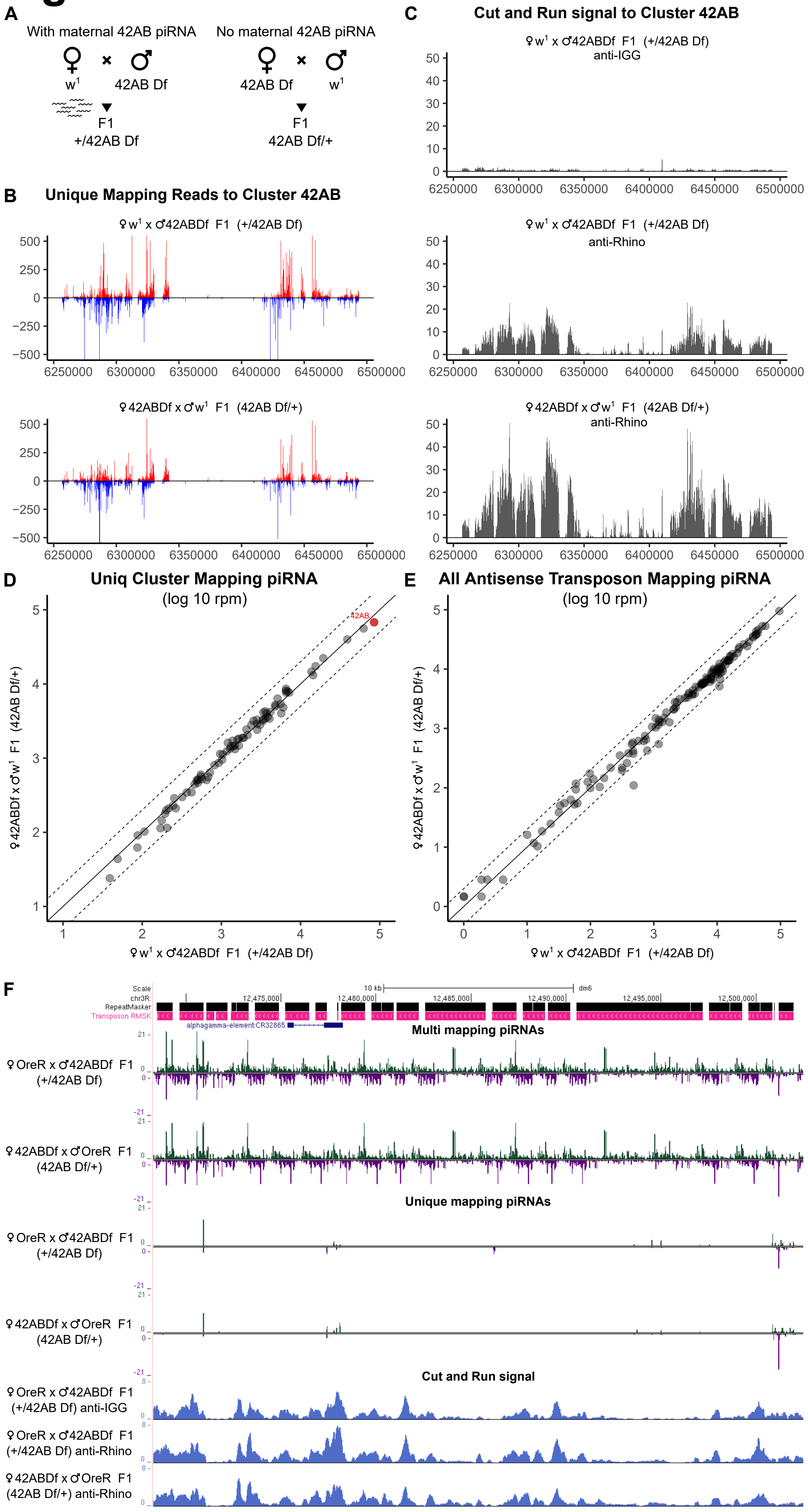
A
With maternal 42AB piRNA
No maternal 42AB piRNA
♀ w1 × ♂ 42AB Df
F1 +/42AB Df
♀ 42AB Df × ♂ w1
F1 42AB Df/+
B Unique Mapping Reads to Cluster 42AB
♀w1 x ♂42ABDf F1 (+/42AB Df)
♀42ABDf x ♂w1 F1 (42AB Df/+)
C Cut and Run signal to Cluster 42AB
♀w1 x ♂42ABDf F1 (+/42AB Df) anti-IGG
♀w1 x ♂42ABDf F1 (+/42AB Df) anti-Rhino
♀42ABDf x ♂w1 F1 (42AB Df/+) anti-Rhino
D Uniq Cluster Mapping piRNA (log 10 rpm)
42AB
♀42ABDf x ♂w1 F1 (42AB Df/+)
♀w1 x ♂42ABDf F1 (+/42AB Df)
E All Antisense Transposon Mapping piRNA (log 10 rpm)
♀42ABDf x ♂w1 F1 (42AB Df/+)
♀w1 x ♂42ABDf F1 (+/42AB Df)
F
Scale
chr3R:
RepeatMasker
Transposon RMSK
10 kb
dm6
12,475,000
12,480,000
12,485,000
12,490,000
12,495,000
12,500,000
alphagamma-element:CR32865
Multi mapping piRNAs
♀OreR x ♂42ABDf F1 (+/42AB Df)
♀42ABDf x ♂OreR F1 (42AB Df/+)
Unique mapping piRNAs
♀OreR x ♂42ABDf F1 (+/42AB Df)
♀42ABDf x ♂OreR F1 (42AB Df/+)
Cut and Run signal
♀OreR x ♂42ABDf F1 (+/42AB Df) anti-IGG
♀OreR x ♂42ABDf F1 (+/42AB Df) anti-Rhino
♀42ABDf x ♂OreR F1 (42AB Df/+) anti-Rhino
